# PHOSPHO1, a novel skeletal regulator of insulin resistance and obesity

**DOI:** 10.1101/2020.05.04.075895

**Authors:** KJ Suchacki, NM Morton, C Vary, C Huesa, MC Yadav, BJ Thomas, S Rajoanah, L Bunger, D Ball, M Barrios-Llerena, AR Guntur, Z Khavandgar, WP Cawthorn, M Ferron, G Karsenty, M Murshed, CJ Rosen, VE MacRae, JL Millán, C Farquharson

**Affiliations:** Roslin Institute, R(D)SVS, University of Edinburgh, Scotland, UK; Centre for Cardiovascular Science, University of Edinburgh, Scotland, UK; Center for Molecular Medicine, Maine Medical Center Research Institute, Scarborough, Maine, USA; MRC Centre for Reproductive Health, University of Edinburgh, Scotland, UK; Sanford Burnham Prebys Medical Discovery Institute, La Jolla, USA; Scottish Rural College, Edinburgh, Scotland, UK; Medical Sciences and Nutrition, School of Medicine, University of Aberdeen, Scotland, UK; Department of Medicine and Faculty of Dentistry, McGill University, Montreal, Canada; Molecular Physiology Research Unit, Institut de recherches cliniques de Montréal, Montreal, Canada; Department of Genetics and Development, Columbia University Medical Center, New York, USA

## Abstract

The skeleton is recognised as a key endocrine regulator of metabolism. Here we show that mice lacking the bone mineralization enzyme PHOSPHO1 (*Phospho1^-/-^*) exhibited improved basal glucose homeostasis and resisted high-fat-diet induced weight gain and diabetes. The metabolic protection in *Phospho1^-/-^* mice was manifested in the absence of altered levels of osteocalcin. Osteoblasts isolated from *Phospho1^-/-^* mice were enriched for genes associated with energy metabolism and diabetes; *Phospho1* both directly and indirectly interacted with genes associated with glucose transport and insulin receptor signalling. Canonical thermogenesis via brown adipose tissue did not underlie the metabolic protection observed in adult *Phospho1^-/-^* mice. However, the decreased serum choline levels in *Phospho1^-/-^* mice were normalized by feeding a 2% choline rich diet resulting in a normalization in insulin sensitivity and fat mass. This study identifies PHOSPHO1 as a potential therapeutic target for the treatment of obesity and diabetes.

## Introduction

In addition to its classical structural functions, the skeleton is a site of significant glucose uptake and is involved in the regulation of whole-body glucose metabolism (1–8). Osteocalcin (OCN) is the most abundant osteoblast-specific non-collagenous protein derived from bone and is thought to maintain the mechanical properties of the bone matrix by regulating calcium binding when fully carboxylated (GLA13-OCN) (9). However, when OCN is not γ-carboxylated (uncarboxylated (GLU-OCN) or undercarboxylated (GLU13-OCN)), it is released from bone into the circulation where it is able to regulate whole-body glucose metabolism in an endocrine manner (7, 10–13). Mice deficient in OCN have increased fat mass and are hyperglycemic, hypoinsulinemic and insulin-resistant in muscle. Furthermore, serum GLU17-OCN (human form of GLU13) levels and β-cell function show an inverse correlation with glycated haemoglobin (HbA1c), fat mass and plasma glucose levels (14–18).

Osteoblasts regulate glucose metabolism through OCN-dependent and independent mechanisms (19, 20). An alternative candidate is the bone-specific cytosolic phosphatase; Phosphatase, Orphan 1 (PHOSPHO1) (21–30). PHOSPHO1 initiates bone matrix mineralization and PHOSPHO1 deficiency causes significant skeletal pathology, bowed long bones, osteomalacia and scoliosis in early life (31–34). In addition to the role of PHOSPHO1 in skeletal biomineralization, PHOSPHO1 has been implicated in the regulation of energy metabolism in humans (35–38). Within the *PHOSPHO1* gene, differential methylation sites have been identified as potentially useful biomarkers for clinical application in the early detection of type-2 diabetes (35) and significant associations between methylation at loci within the *PHOSPHO1* gene and the future risk of type-2 diabetes exist (36, 37). Differential methylation in *PHOSPHO1* was associated with three lipid traits (total cholesterol, high-density lipoprotein cholesterol, and triglycerides) (39). Most recently, genetic variants of *PHOSPHO1* in a bivariate twin study were found to be associated with body mass index and waist-hip ratio (38). Taken together these findings suggest that in addition to the established role of PHOSPHO1 in biomineralization of the skeleton and dentition, *Phospho1* ablation may result in improved glucose homeostasis and a reduction in metabolic disease susceptibility. To address this, we examined the metabolic phenotype of juvenile and adult *Phospho1^-/-^* mice.

## Results

### *Phospho1* inactivation improves glucose tolerance and insulin sensitivity in juvenile mice

Growth of *Phospho1^-/-^* mice was decreased compared to wild-type (WT) mice (Figure 1a). Juvenile *Phospho1^-/-^* mice (35-day-old) had reduced body weight and blood glucose levels (WT: 9.31±0.31 mmol/L, *Phospho1^-/-^*: 7.76±0.32 mmol/L; p<0.01) (Figures 1b & c), improved glucose tolerance (Figure 1d) and whole body insulin sensitivity compared to WT counterparts (Figure 1e). Consistent with this, adipose depots were smaller in *Phospho1^-/-^* mice: inguinal (iWAT; WT: 4.31±0.27 mg/g, *Phospho1^-/-^*: 2.58±0.20 mg/g; p<0.001), mesenteric (mWAT; WT: 5.30±0.30 mg/g, *Phospho1^-/-^*: 3.39±0.40 mg/g; p<0.01) and gonadal (gWAT; WT: 4.31±0.27 mg/g, *Phospho1^-/-^*: 2.58±0.20 mg/g; p<0.001) adipose tissue (Figure 1f). *Phospho1^-/-^* mice had significantly smaller livers (WT: 64.36±0.49 mg/g, *Phospho1^-/-^*: 52.71±3.37 mg/g; p<0.05), and quadriceps (WT: 6.39±0.36 mg/g, *Phospho1^-/-^*: 5.01±0.26 mg/g; p<0.01) (Figure 1g). Food intake (WT: 0.13±0.01 g/gBW/day; *Phospho1^-/-^*: 0.12± 0.01 g/gBW/day) (Figure 1i), activity (Supplementary data 1) and energy expenditure (day and night respiratory exchange rate (RER)) (Figure 1j) were comparable between genotypes.

**Figure 1.**
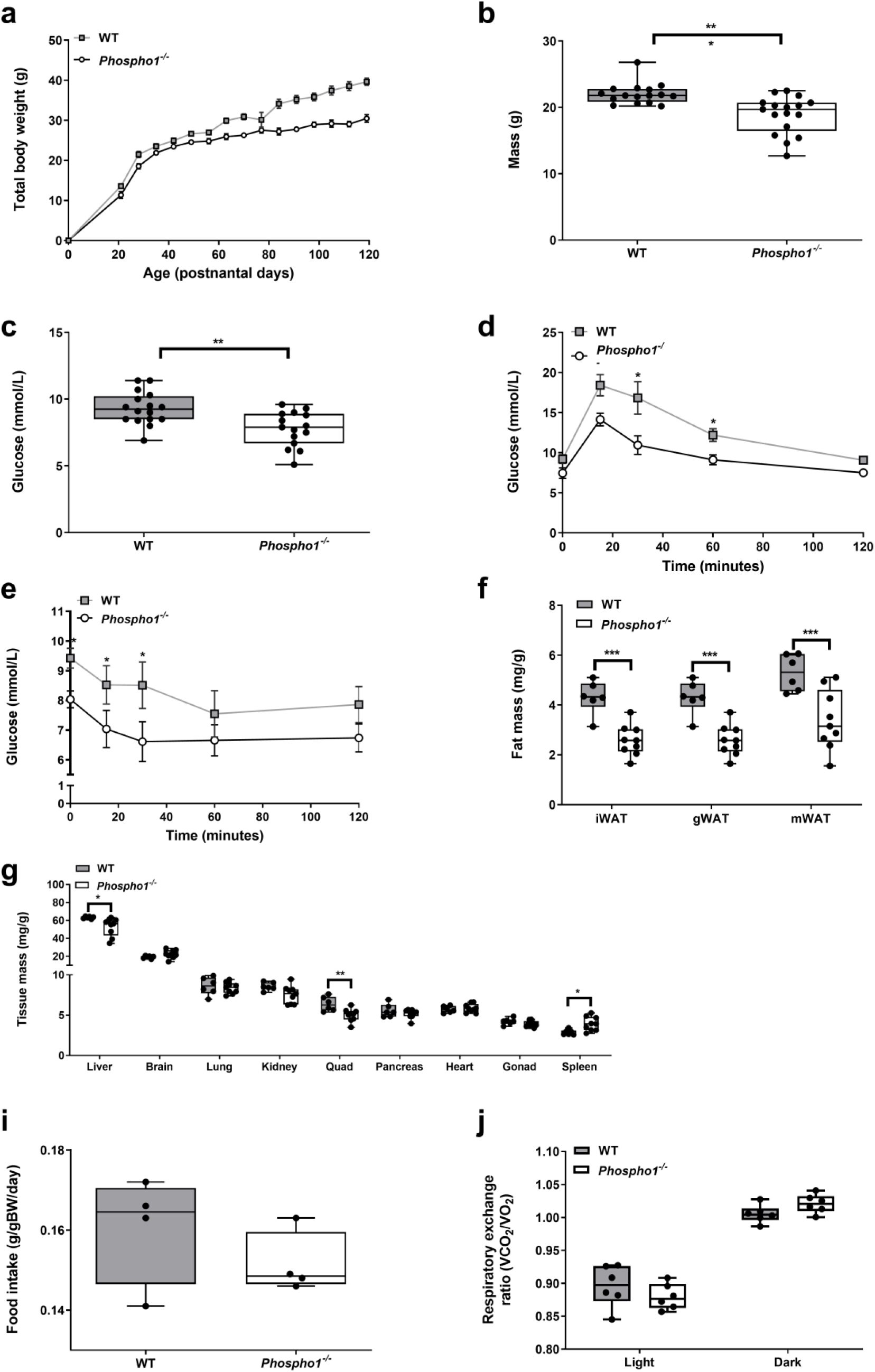
Juvenile *Phospho1^-/-^* mice display increased insulin sensitivity and decreased fat mass. *Phospho1^-/-^* mice showed decreased growth and live weight at 35days of age **(a-b)** and fasting glucose **(c),** improved glucose and insulin tolerance **(d-e)** and decreased adipose tissue **(f).** Notable differences in tissue mass were also observed in the liver, quadriceps and spleen. These changes were not a consequence of altered food intake **(i)** or energy expenditure **(j)**. Data are represented as mean ±S.E.M. *p<0.05, **p<0.01, ***p<0.001.

### *Phospho1* deficiency protects from diet-induced diabetes in adult mice

We next fed WT and *Phospho1^-/-^* mice a chronic high fat diet (HFD) from weaning until adulthood (120 days of age). Adult *Phospho1^-/-^* mice maintained lower body weight with HFD but did not resist weight gain (Control diet (CD) - WT: 34.20 ±1.12g, *Phospho1^-/-^*. 28.30±0.59g; HFD-WT: 38.0±1.54 g, *Phospho1^-/-^*. 32.4±1.26 g; p<0.05; Figure 2a)). Fasting glucose levels were raised in WT mice fed a HFD but not in *Phospho1^-/-^* mice (CD-WT: 9.50 ±0.37mmol/l, *Phospho1^-/-^*: 8.59±0.27mmol/l; HFD-WT 10.3±0.53mmol/l, *Phospho1^-/-^*: 9.27±0.77) (Supplementary data 2).

**Figure 2.**
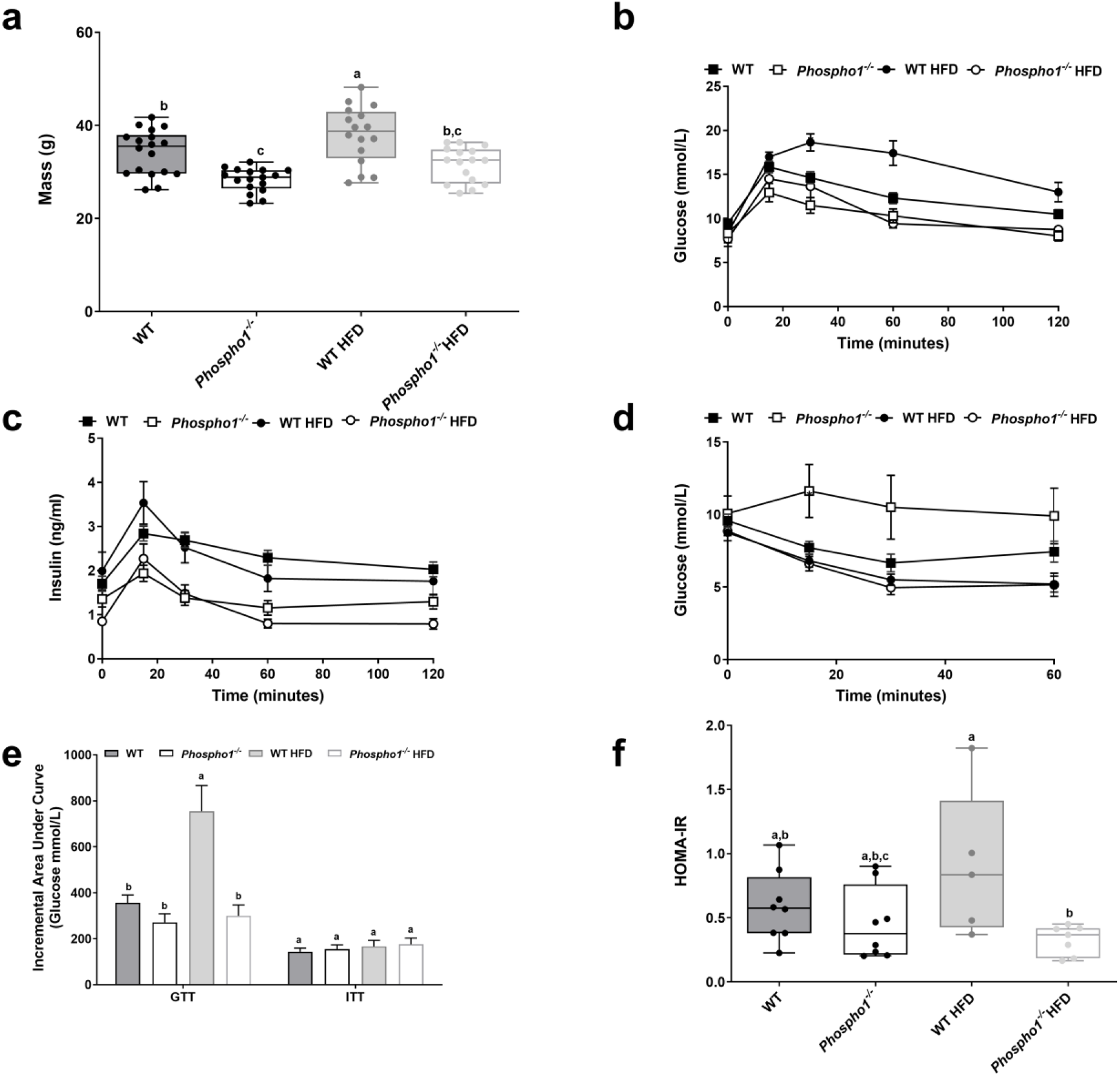
*Phospho1^-/-^* mice are protected from glucose intolerance. **(a)** Body mass **(b)** glucose tolerance test (GTT) **(c)** glucose stimulate insulin secretion (GSIS) **(c)** insulin tolerance test (ITT) **(e)** incremental area under the curve for GTT and ITT, **(g)** HOMA-IR. Different letters above the error bar for each gene show significant difference at p<0.05.

Glucose tolerance was improved in *Phospho1^-/-^* mice after chronic HFD (Figure 2b). Insulin secretion across the glucose tolerance test (GTT) was also lower in *Phospho1^-/-^* mice on both CD and HFD, suggestive of insulin sensitisation rather than exaggerated β-cell insulin secretion as the major basis of the phenotype (Figure 2c). This was confirmed with insulin tolerance tests (ITT) after chronic HFD, which revealed greater glucose disposal in *Phospho1^-/-^* mice (Figure 2d-f).

### *Phospho1* deficiency protects from diet-induced obesity in adult mice

The insulin sensitivity observed in *Phospho1^-/-^* mice was consistent with the finding of smaller inguinal (CD-WT: 4.51±0.37 mg/g BW, *Phosphol^-/-^*. 2.79±0.42 mg/g BW; HFD-WT: 14.67 ±2.12 mg/g BW, *Phospho1^-/-^*7.95 ±1.56 mg/g BW; p<0.01), mesenteric (CD-WT: 13.2±1.34 mg/g BW, *Phosphol^-/-^*. 5.56±1.61 mg/g BW; HFD-WT:24.14 ±4.05 mg/g BW, *Phospho1^-/-^*:10.22 ±1.57 mg/g BW; p<0.01) and gonadal (CD-WT: 13.7±1.81 mg/g BW, *Phospho1^-/-^*: 6.96±0.58 mg/g BW; HFD-WT:28.77 ±3.12 mg/g BW, *Phosphor^-/-^*:18.78 ±2.37 mg/g BW; p<0.01) fat depots noted in CD and HFD *Phospho1^-/-^* mice at necropsy (Figure 3a). Moreover, confirmation that *Phospho1^-/-^* mice did not become obese when fed a HFD, was shown by μMRI and CT (Supplementary data 3). These observations were also not explained by altered activity or increased food intake in 120-day-old adult male mice (data not shown).

**Figure 3.**
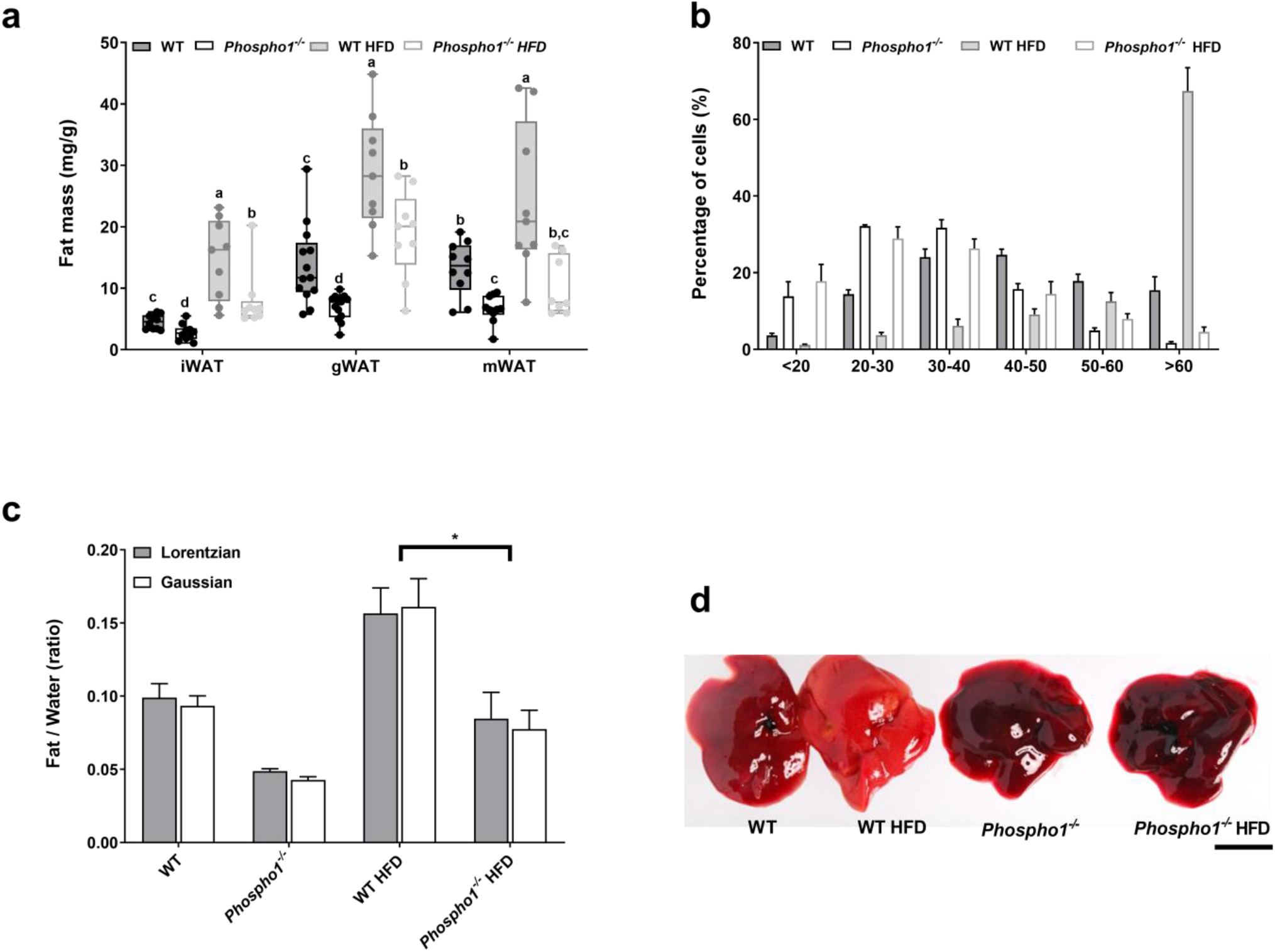
*Phospho1^-/-^* are protected from NAFLD. **(a)** Fat analysis of 120-day-old WT and *Phospho1^-/-^* mice on both a control and HFD. **(b)** Quantification of gonadal fat adipocyte diameter **(c)** Quantitative assessment of liver fat utilising spectroscopy. **(d)** Gross livers of representative mice left to right (WT, WT HFD, *Phospho1^-/-^, Phospho1^-/-^* HFD; scale bar = 10mm) Data are represented as mean ±S.E.M. *p<0.05, **p<0.01, ***p<0.001. Different letters above the error bar for each gene show significant difference at p<0.05.

Histological analysis revealed smaller gonadal adipocytes in *Phospho1^-/-^* mice fed both a CD and a HFD. High fat feeding had no significant effect on gonadal adipocyte size in *Phospho1^-/-^* mice however it significantly increased the number of large adipocytes (> 60 μm in diameter) in WT mice (p<0.0001) (Figure 3b). *Phospho1^-/-^* mice were also protected from the pronounced hepatic fat accumulation that was noted in WT mice following HFD feeding (Figures 3c & d).

### Insulin sensitivity in *Phospho1^-/-^* mice is independent of elevated adiponectin serum levels

In an attempt to uncover the endocrine mechanism(s) responsible for the increased insulin sensitivity in *Phospho1^-/-^* mice, serum levels of adiponectin and leptin were measured. Levels of high molecular weight adiponectin, a hormone linked to insulin-sensitisation (40), were decreased in *Phospho1^-/-^* mice fed either a CD (2.62-fold) or a HFD (1.92-fold) (both p<0.001) suggesting that insulin sensitivity and protection from obesity are independent of adiponectin (Figure 4a). The observed decrease in circulating adiponectin in *Phospho1^-/-^* mice was not due to decreased bone marrow adipose tissue, an endocrine organ that contributes significantly to serum adiponectin (Supplementary data 4) (41).

**Figure 4.**
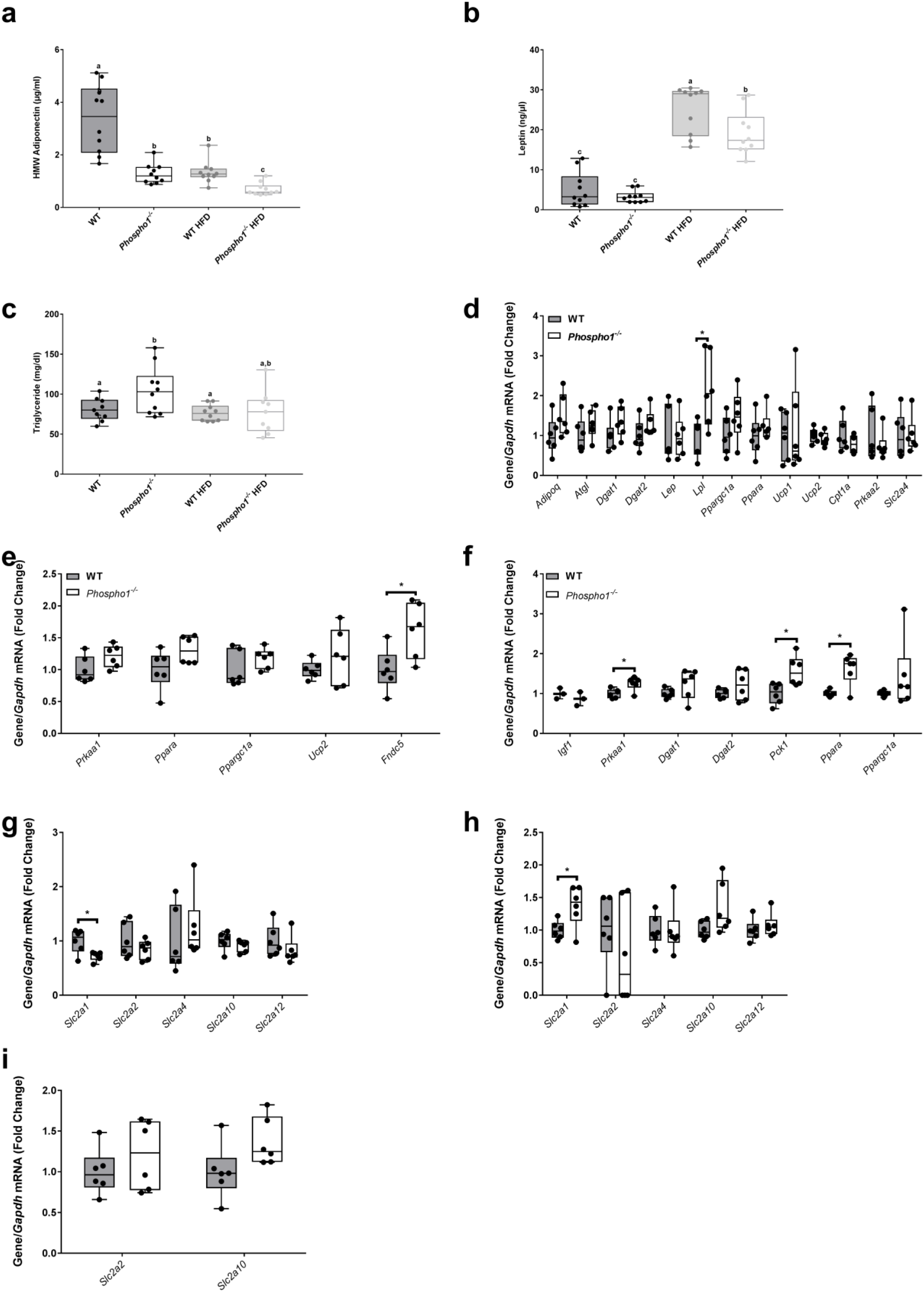
*Phospho1^-/-^* mice are insulin sensitive despite decreased adiponectin. **(a)** Adiponectin, **(b)** leptin and **(c)** triglyceride serum quantification. RT-qPCR analysis of tissue extracted from 120 day old WT and *Phospho1^-/-^* mice **(d)** Adipose tissue **(e)** Quadriceps Femoris and **(f)** Liver. RT-qPCR analysis of GLUT receptors from **(g)** Adipose tissue **(h)** Quadriceps Femoris and **(i)** Liver. Data are represented as mean ±S.E.M. * p<0.05, **p<0.01, ***p<0.001. Different letters above the error bar for each gene show significant difference at p<0.05.

Serum leptin levels in CD fed mice were unaffected by *Phospho1* -deficiency, whereas in comparison to WT mice fed a HFD serum leptin levels were significantly decreased 1.31-fold (p<0.05) in *Phospho1^-/-^* mice fed a HFD (Figure 4b), accordant with reduced fat mass (42). *Phospho1^-/-^* CD mice had increased circulating serum triglycerides compared to WT CD mice, but no change was observed in WT and *Phospho1^-/-^* mice fed a HFD (Figure 4c). Consistent with increased oxidative metabolism of carbohydrate and lipids in other peripheral tissues, mRNA levels of genes encoding key metabolic proteins were increased in adipose tissue *(Lpl)* muscle *(Fndc5)* and liver *(Prkaa1, Pepck1* and *Ppara)* (Figures 4d-f). The mRNA levels of genes encoding GLUT receptors (Slc2a1, 2, 4, 10 and 12) were largely unchanged (Figure 4g-i).

### Canonical thermogenesis does not underlie the metabolic protection observed in adult *Phospho1* deficient mice

35 day *old-Phospho1^-/-^* mice had decreased interscapular brown adipose tissue (BAT) mass compared to WT counterparts (WT: 5.11±0.57 mg/g, *Phospho1^-/-^*: 3.05±0.40 mg/g; p<0.01) (Figure 5a). However this reduction in BAT mass did not persist to adulthood nor during high-fat feeding (Figure 5b). Strikingly, adult and high-fat-fed *Phospho1^-/-^* mice had smaller brown adipocytes compared to WT controls (Figure 5c). In order to see if BAT activation and thermogenesis might be responsible for the observed phenotype we measured key brown fat genes including uncoupling protein 1 *(Ucp1)* (Figures 5d & e), no differences were observed in the mRNA and protein levels. Furthermore, there were no significant differences in respiratory exchange ratio (RER, indicative of metabolic substrate preference) or energy expenditure between WT or *Phospho1^-/-^* mice fed either a chow or HFD housed at either room temperature or during cold exposure (4°C) (Figures 5f-i). These *in vivo* data show that increased canonical thermogenesis does not underlie the metabolic protection observed in the *Phospho1* deficient mice so this area was not pursued further.

**Figure 5.**
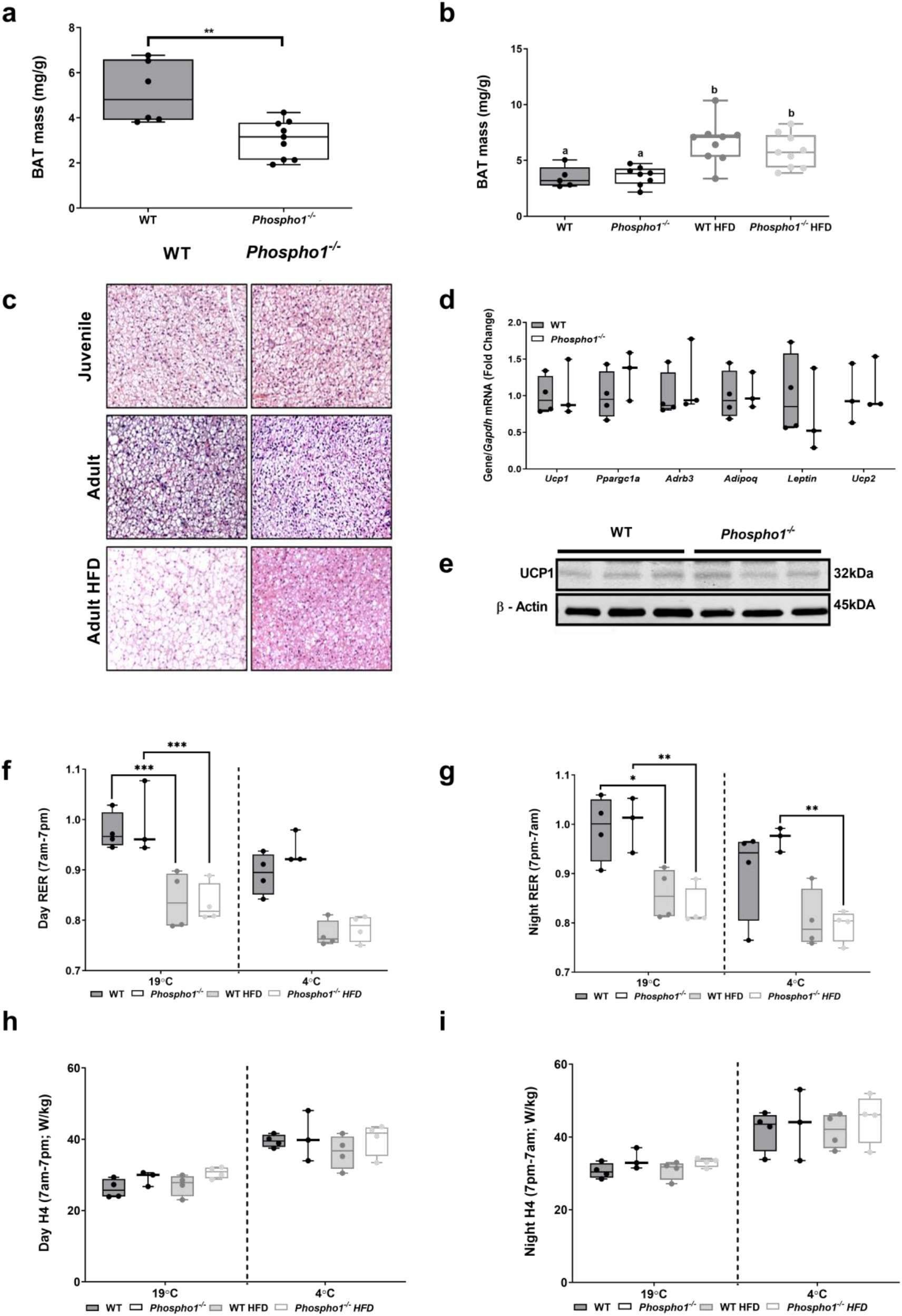
Canonical thermogenesis does not underlie the metabolic protection observed in the adult *Phospho1* deficient mice. Brown adipose tissue (BAT) mass in juvenile (35 day old; **a**) and adult (120 day old) WT and *Phospho1^-/-^* mice **(b).** Representative micrographs of BAT from WT and *Phospho1^-/-^* mice **(c). (d)** Brown fat gene expression and **(e)** UCP1 protein analysis. Insulin sensitivity and protection from diet induced obesity in *Phospho1^-/-^* mice was not a consequence of altered energy expenditure (RER - respiratory exchange ratio; H4 = H3 (W) / Lean mass (Kg)) **(f-i).** Data are represented as mean ±S.E.M. *p<0.05, **p<0.01, ***p<0.001. Different letters above the error bar for each gene show significant difference at p<0.05.

### Diabetes mellitus-associated genes are enriched in *Phospho1^-/-^* primary osteoblasts

We next sought to unravel the genetic circuitry responsible for the improved glucose tolerance in *Phospho1^-/-^* mice. To address this, a transcriptomic analysis of *Phospho1* deficient osteoblasts was conducted. There was a striking 20-fold up-regulation of embryonic stem cell phosphatase *(Esp)* mRNA, the gene encoding the protein osteotesticular protein tyrosine phosphatase (OST-PTP) in *Phospho1* deficient mice (7). These data were validated in primary WT and *Phospho1^-/-^* calvarial osteoblasts and *Phospho1* deficient osteoblast overexpressing *Phospho1* (Figures 6a-b). Protein tyrosine phosphatases are recognised master regulators of insulin receptor signalling (INSR), negatively modifying osteoblast-insulin signalling and thereby controlling GLU13-OCN release (7, 43–47). The identification of elevated *Esp* expression in *Phospho1^-/-^* mice was strongly suggestive of a reciprocal regulation between OST-PTP and PHOSPHO1 in the control of glucose homeostasis. This increased *Esp* expression was however inconsistent with the improved glucose tolerance in the *Phospho1^-/-^* mice. Therefore in an attempt to reconcile this anomaly we measured circulating GLU–OCN and GLU13-OCN, which were found to be unchanged in juvenile and adult *Phosphol^-/-^* mice (Figure 6c). These data implied that elevated serum levels of uncarboxylated or undercarboxylated OCN did not mediate the improved metabolic phenotype (Figures 6c & d). The elevated levels of carboxylated and total OCN in *Phospho1* deficiency was consistent with increased bone turnover in these mice as previously reported (31).

**Figure 6.**
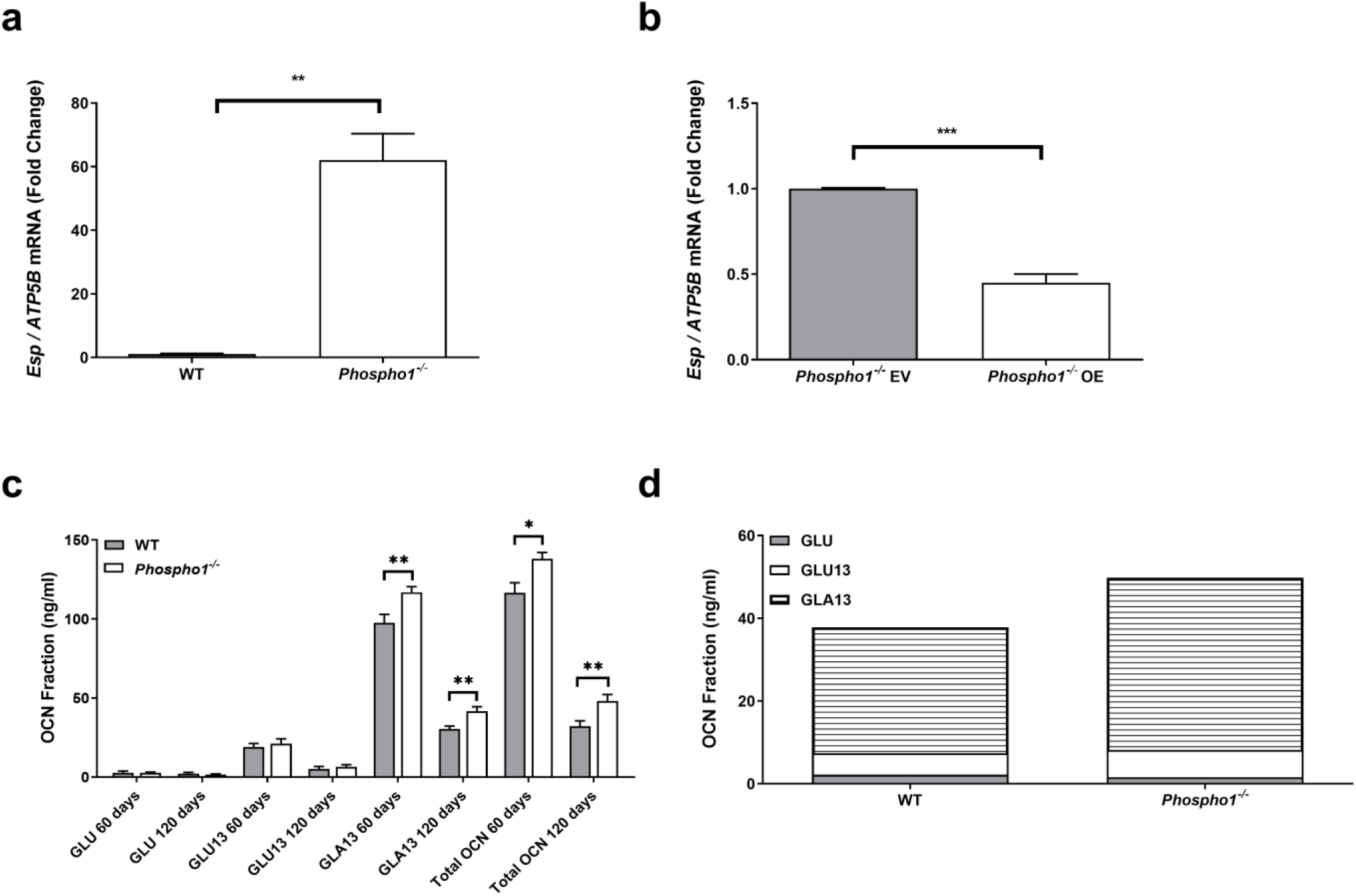
Osteocalcin-independent mechanism of PHOSPHO1-regulated energy metabolism. To assess the relative change in *Esp* mRNA expression in primary calvarial osteoblasts, RT-qPCR was conducted to compare **(a)** primary WT and *Phospho1^-/-^* osteoblasts and **(b)** *Phospho1^-/-^* osteoblast transfected with empty (EV) or overexpressing (OE) vectors. Osteocalcin content of serum from WT and *Phospho1^-/-^* mice at 60 and 120 days of age **(c,d)**. Data are represented as mean ±S.E.M. * p<0.05, **p<0.01, ***p<0.001.

### *Phospho1^-/-^* calvarial osteoblasts show a greater metabolic capacity in utilizing exogenous substrates compared to WT osteoblasts

Primary calvarial osteoblast metabolism analysis revealed that differentiated *Phospho1^-/-^* osteoblasts had elevated basal oxygen consumption rates (indicative of oxidative phosphorylation) compared to WT osteoblasts when supplied exogenously with glucose, pyruvate and glutamine (Supplementary data 5). Following inhibition of complex V of the electron transport chain using oligomycin, cells upregulated glycolytic rates which is estimated from extracellular acidification rate (ECAR). There was a significant increase in the glycolytic rate of differentiated *Phospho1^-/-^* osteoblasts compared to WT osteoblasts suggesting that they have increased glucose metabolism. A recent study (48) showed that CO_2_ production from oxygen consumption rate (OCR) caused changes in the acidification rates. We corrected for this by calculating the acidification rate from CO_2_ and subtracting it from the total acidification rate to obtain the glycolytic acidification rates. We showed the glycolytic rates (proton production rates, PPRGlyc) were altered after oligomycin treatment (Supplementary data 5). In sum, differentiating *Phospho1^-/-^* calvarial osteoblasts show a greater metabolic capacity in utilizing exogenous substrates compared to WT osteoblasts.

To further clarify the genetic pathways underpinning these observations, 22 differentially expressed genes identified from the microarray analysis were found by Ingenuity Pathway Analysis (IPA) to be associated with glucose homeostasis (Supplementary data 6). Further predictive analysis of the differentially expressed genes of the original microarray data set identified > 40 genes to be associated with energy metabolism (p=1.04×10^-6^) (Figure 7a) of which 10 were found to be associated with both diabetes and bone, following an NCBI (PubMed) *in silico* search. Validation by RT-qPCR confirmed that all 10 genes (*Vdr, Mpeg1, Slc1a3, Adamts4, Bmp4, Cd68, Cfp, Cxcl4, Fmod and Lum*) were differentially regulated in *Phospho1* deficient osteoblasts as predicted by IPA (p<0.05; Figure 7b & Supplementary data 7). Furthermore, GeneMANIA network analysis predicted that *Phospho1* both directly and indirectly interacts with 36 genes associated with glucose transport and metabolic processes and insulin receptor signalling (Figure 8). Of the output genes, *Atf4, Foxo1* and *Insr* are recognized skeletal modulators of energy metabolism, suggesting crosstalk between *Phospho1* and other metabolic regulatory genes (5, 49, 50).

**Figure 7.**
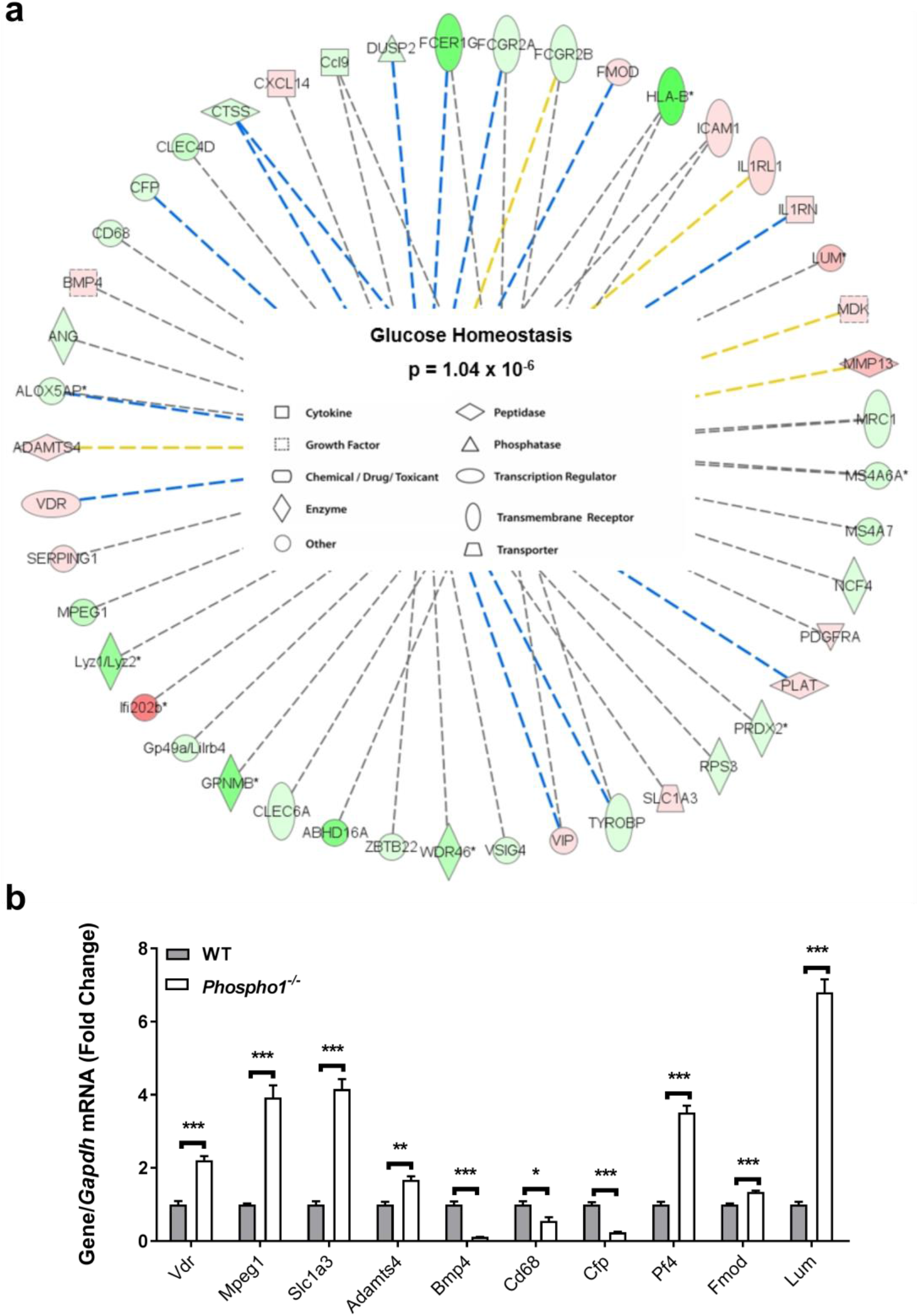
Ingenuity Pathways Analysis network summary predictions. Ingenuity Pathways Analysis was used to predict further genes associated with glucose homeostasis based upon the 22 genes found to be differentially expressed in the microarray **(a)**. **(b)** Predicated genes were confirmed by RT-qPCR in WT and *Phospho1^-/-^* primary calvarial osteoblast’s. Results were normalized to the *Atp5b* housekeeping gene. Data are represented as mean ±S.E.M (n=3 replicates). * P<0.05, ** P<0.01, ***P<0.001. Red = Up regulated. Green = Down regulated (the darker the shade of green and red colour indicates a more extreme up/down regulation, conversely the paler the shade indicates a more subtle up/down regulation. Dashed line = indirect interaction (blue = inhibition, yellow = findings underlying the relationship are inconsistent with the state of the downstream node, grey = Ingenuity Pathways Analysis prediction).

**Figure 8.**
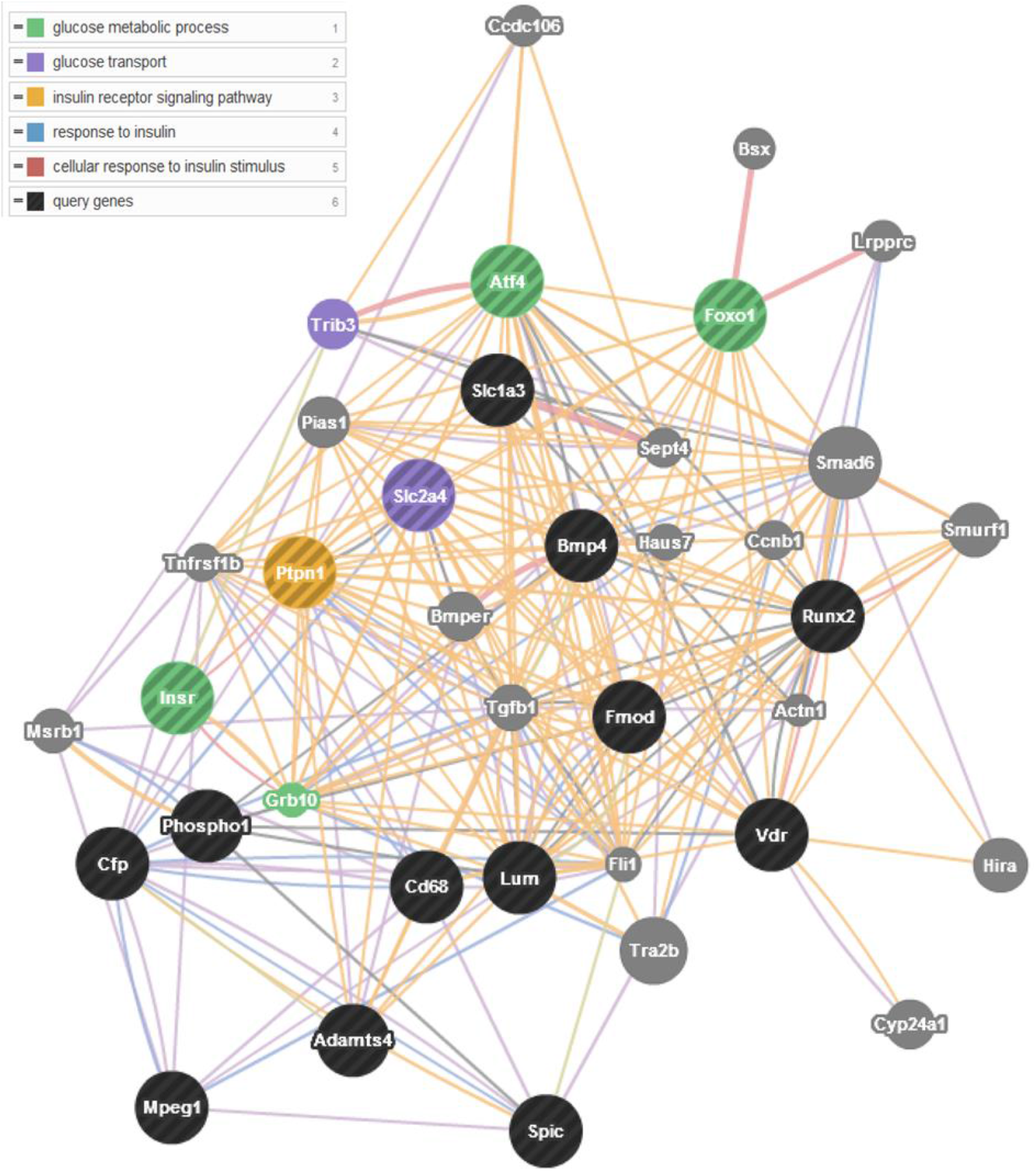
GeneMANIA network summary predictions. GeneMANIA network generated using Ingenuity Pathways Analysis gene predictions. The network highlights potential interactions between *Phospho1* and related osteoblast genes involved in the glucose metabolic process, encompassing; glucose transport, insulin receptor signalling, response to insulin and cellular response to insulin stimulus. Query genes (black) with the exception of Spic and Runx2 which were inputted manually, other genes (grey) were generated by the programme using a large set of inbuilt functional association data. Node size are based on GO terms. Network line colour corresponds to interaction: purple = co-expression, pink = physical interactions, blue = co-localisation, green = shared protein domains orange = predicted, grey = other

### Identification of differentially expressed serum proteins in *Phospho1^-/-^* mice

To explore the secretome profile, quantitative SWATH (sequential window acquisition of all theoretical spectra) MS (mass spectrometry) proteomics (51) was conducted on serum from WT and *Phospho1^-/-^* mice fed CD and HFD. Differentially expressed proteins (> 100) were identified in HFD *Phospho1^-/-^* serum compared to HFD-fed WT mice. These proteins were highly associated with glycolysis, gluconeogenesis and ‘metabolic pathways’ (Supplementary data 8). Pathway and network analysis predicted that the identified proteins interacted with miR-34a; a microRNA that is known to affect diverse parts of insulin signalling in the pancreas, liver, muscle and adipose tissue (52).

### Bone-derived choline is involved in global energy regulation

Neutral sphingomyelinase 2 (nSMase2) catalyses the hydrolysis of sphingomyelin to form ceramide and phosphocholine (PCho) (53). Furthermore, PCho is the preferred substrate for PHOSPHO1 yielding choline and inorganic phosphate (Pi) (Figure 9a) (54). As elevated levels of both ceramide and choline result in insulin resistance in mice (55, 56) we targeted these intermediates to establish if alterations in their serum levels could explain the insulin sensitive phenotype in *Phospho1^-/-^* mice (55, 56). The levels of various ceramide species were unchanged (Figure 9b) however *Phospho1^-/-^* mice had a significant decrease in serum choline levels (WT: 0.152±0.001 μg/ml, *Phospho1^-/-^*: 0.128±0.003 μg/ml; p<0.01) (Figure 9c) which were normalized upon choline supplementation. Supplementation of WT and *Phospho1^-/-^* mice with a 2% choline diet (a well-tolerated, palatable diet (57)), also normalized the insulin sensitivity measured in *Phospho1^-/-^*, measured by GTT (Figure 9d). However, unlike WT mice which when fed a 2% choline diet took longer to recover from the insulin challenge, *Phospho1^-/-^* showed no metabolic change in response to insulin between the diets (Figure 9e). Furthermore, choline supplementation normalized the lean phenotype observed in *Phospho1^-/-^* mice (Figure 9f & g). These results support the notion that *Phospho1* -deficiency improves the metabolic profile of mice *in vivo* and confers resistance to obesity and diabetes in part via the alteration of serum choline levels.

**Figure 9.**
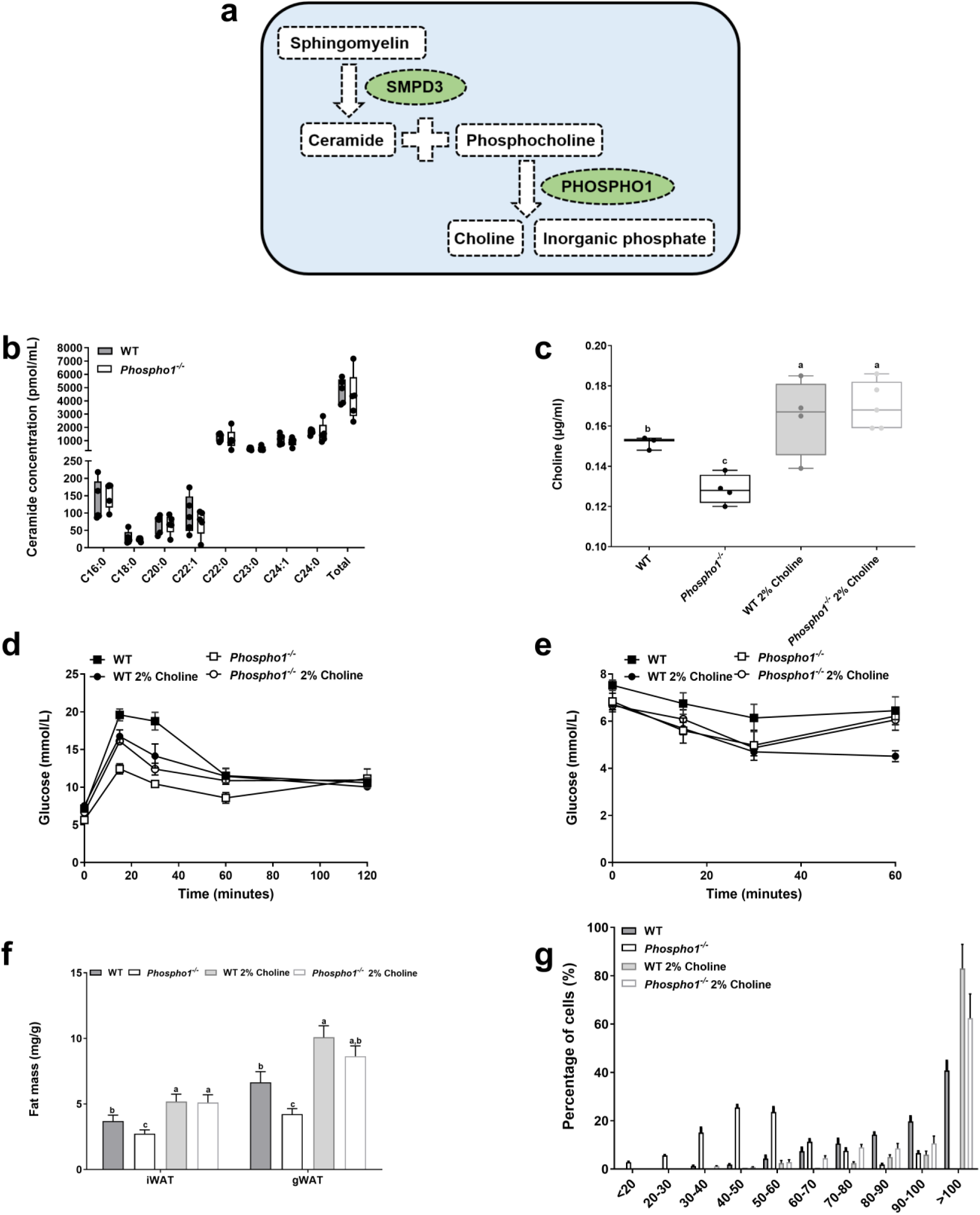
Bone derived choline regulates insulin sensitivity. **(a)** Schematic diagram outlining the mechanisms by which ceramide and choline and linked. **(b)** Mouse serum ceramide and choline **(c)** analysis by LC-MS/MS (ceramide) and assay (choline). **(d)** GTT **(e)** ITT **(f)** Dissected fat depot weights **(g)** quantification of epididymal fat adipocyte diameter and representative histology. Data are represented as mean ±S.E.M. *p<0.05, **p<0.01, ***p<0.001. Different letters above the error bar for each gene show significant difference at p<0.05.

## Discussion

The fundamental observations presented in this study further strengthen the concept that bone acts as an endocrine organ and provides empirical evidence for a critical role for PHOSPHO1 in energy metabolism. *Phospho1-deficiency* results in decreased blood glucose levels, improved insulin sensitivity, glucose tolerance and conferred protection from diet-induced obesity and diabetes in mice despite a 60-fold up-regulation of *Esp* expression by *Phospho1^-/-^* osteoblasts. Mice lacking *Esp* in osteoblasts present with severe hypoglycemia and hyperinsulinemia resulting in postnatal lethality in the first two weeks of life (7). Conversely, mice overexpressing *Esp* exclusively in osteoblasts were glucose intolerant and insulin resistant (7, 43–47). Intriguinly, the increased insulin sensitivity in *Phospho1* mice was not associated with the expected rise in serum GLU13-OCN levels suggesting that PHOSPHO1-regulated energy metabolism is via OCN-independent mechanisms. This notion has previously been observed when partial genetic ablation of osteoblasts profoundly affected energy expenditure, gonadal fat weight and insulin sensitivity which were not restored by the administration of OCN (19, 20). Nevertheless, it is possible that the increased insulin sensitivity noted in the *Phospho1^-/-^* mice may be primed by an initial rise in GLU13-OCN levels, which is eventually normalized in a compensatory manner by the observed increase in *Esp* expression. This being the case we would predict that the loss of *Esp* on a *Phospho1^-/-^* background would exacerbate the insulin sensitivity due to increased GLU13-OCN serum levels. These data strengthen the concept that a novel pathway exists between osteoblasts and glucose homeostasis, however it does highlight the potential cross-talk between OCN-dependent and OCN-independent mechanisms of glucose metabolism.

The role of PHOSPHO 1 in controlling bone mineralization has been extensively investigated through the use of both *in vitro* and *in vivo* mouse models. Crucial for the initiation of mineralization within matrix vesicles, PHOSPHO1 hydrolyzes membrane lipid derivatives, primarily PCho to produce Pi (utilized in hydroxyapatite formation) and choline (54). Phosphocholine can be formed from choline via choline kinase activity or phosphatidylcholine via PLA2 and ENPP6 as well as from the hydrolysis of sphingomyelin, via nSMase2 to form PCho and ceramide (58). Mindful of this, it has been reported that elevated levels of both ceramide and choline result in insulin resistance in mice (55, 56). We saw no change in ceramide species in *Phospho1^-/-^* mice, however there was a significant decrease in serum choline levels in *Phospho1^-/-^* mice, which was normalized in *Phospho1^-/-^* mice fed a 2% choline rich diet resulting in a normalisation in insulin sensitivity and fat mass. This study highlights for the first time the importance of bone-derived choline in the regulation of energy metabolism, however it remains clear that choline supplementation alone does not fully correct all metabolic defects in *Phospho1^-/-^* mice.

The regulation of global energy metabolism by the skeleton is a complex, multifactorial process, and it is therefore likely that, as yet, undefined bone secreted factors may play a significant role in PHOSPHO1 regulated global energy regulation. We identified >100 secreted proteins, that may also be involved in the energy regulation via the skeleton. These unique proteins were highly associated with glycolysis, gluconeogenesis and ‘metabolic pathways’ and association with miR-34a; a microRNA that affects diverse parts of insulin signalling in the pancreas, liver, muscle and adipose tissue (52, 59). Furthermore, lumican, a proteoglycan secreted by differentiating osteoblasts and a constituent of the bone matrix was found to be enriched in serum of *Phospho1^-/-^* mice by both proteomic and microarray analysis. Interestingly, lumican has also been observed in the decidua of diabetic patients (60, 61). Further investigations of these and other candidates may uncover new skeletal regulators of energy metabolism.

The metabolic phenotype observed in the *Phospho1* deficient mice was not due to increased BAT activation despite striking histological differences between WT and *Phospho1^-/-^* tissue. BAT is a thermogenic organ that increases energy expenditure to generate heat, maintaining body temperature in a cold environment (62). When activated by cold exposure or through activation of β3-adrenergic receptors, BAT improves insulin sensitivity and lipid clearance, highlighting its key role in metabolic health (63). PHOSPHO1 has been previously suggested to have a role in murine BAT function. Increased expression of *Phospho1* in both isolated brown and white adipocytes was observed following treatment with the β-adrenergic agonist (CL316,243) compared to placebo treated white adipocytes (64). Mice with an adipose-specific defect in fatty acid oxidation (*Cpt2A^-/-^*) showed a loss of β3-adrenergic induced *Phospho1* expression in BAT (64) and expression of *Phospho1* was shown to be elevated in *Ucp1*-deficient animals; however WT mice had very low *Phospho1* expression in BAT and no-detectable PHOSPHO1 was observed in WT iWAT and BAT (65). No difference in *Phospho1* gene expression was observed between human supraclavicular and subcutaneous adipose progenitor cells (GEO Dataset: GDS5171/8016540) and murine WAT had elevated *Phospho1* gene expression compared to BAT (GEO Dataset: GDS2813/1452485_at). In our hands, we have noted high expression of *Phospho1* in murine BAT but no protein expression was detected (supplementary data 9). Taken together, these data suggest that *Phospho1* is likely to play a role in BAT function, however the BAT phenotype observed in *Phospho1* deficient mice does not appear to underlay the metabolic protection we see in these animals. Further studies are necessary to unravel the role of PHOSPHO1 in BAT using a BAT conditional knock-out model of *Phospho1.*

Collectively, the results of this study add further credibility to the concept that GLU13-OCN is not the sole mediator of the endocrine function of the skeleton (19). We suggest, as others have, that further bone-derived proteins/lipids work in partnership with OCN to regulate the metabolic function of the skeleton (19). Indeed, this study has identified other potential protein mediators and raised the possibility that bone derived choline may contribute to the regulation of the development of the metabolic syndrome. Furthermore, our results suggest that *Esp* may act as a fine controller of insulin sensitivity in mice, offering protection from severe hypoglycaemia and dyslipidaemia.

Several previous reports have suggested an association between PHOSPHO1 expression in disorders of altered energy metabolism such as obesity and diabetes (35–39). The data from this present study is both supportive of such an association but also provides insight into the mechanisms by which PHOSPHO1 may contribute to the regulation of energy metabolism, inclusive of insulin sensitivity, glucose tolerance and fat metabolism. Also, inhibitors of PHOSPHO1 activity such as the proton pump inhibitor lansoprazole, commonly prescribed to control and prevent symptoms of gastroesophageal reflux disease and dyspepsia have been associated with improved glycaemic control in diabetic patients (29, 66–68). Taken together, the identification of PHOSPHO1 in the role of energy metabolism in both the human and mouse offers the potential to manipulate key targets of the PHOSPHO1 pathway to improve metabolic health (36, 37).

## Methods

### Reagents

All chemicals, tissue culture medium and buffers were from Sigma-Aldrich (Dorset, UK) and Invitrogen (Paisley, UK) unless otherwise stated. PCR oligonucleotides were purchased from MWG Eurofins (Ebersberg, Germany) and Primer Design (Southampton, UK). PHOSPHO1 HuCAL Fab bivalent antibody was purchased from AbD Serotech (Kidlington, UK). All antibodies were diluted 1:1,000 unless otherwise noted.

### Animals

*Phospho1* null mice were generated as previously described (32). Offspring carrying the mutant *Phospho1* gene were identified by genotyping (F: 5’-TCCTCCTCACCTTCGACTTC-3’, R: 5’-TCCTCCTCACCTTCGACTTC-3’). All *in vivo* studies were conducted at 120 days of age unless otherwise stated. Male mice were fed a high fat diet consisting of 58% of calories from fat (DBM Scotland, Broxburn, UK) or control diet (6.2% calories from fat; Harlan Laboratories, Indianapolis, IN, USA) starting at 4 weeks of age. Male mice were fed a 2% supplemented choline diet (Harlan Laboratories) or control diet (Harlan Laboratories) for 5 weeks prior to cull at 120 days. Ad libitum food consumption was monitored for 6 days and basal nocturnal activity was quantified using an AM524 Single Layer X, Y IR activity monitor and associated Amonlite software (Linton Instrumentations, Norfolk, UK). Juvenile metabolic activity was measured using indirect calorimetry (Oxymax Lab Animal Monitoring System: CLAMS (Columbus Instruments, OH USA). Adult metabolic rate was measured using indirect calorimetry (TSE PhenoMaster 1.0, with software version 6.1.9). Cold exposed mice were first housed in these cages for 3 days at room temperature (RT) for acclimation and baseline measurements. Mice were then house for 72 hours at 4°C. All experiments were conducted blind to the operator. Animals were maintained under conventional housing conditions with a 12 hour light/dark cycle with free access to food and water (except when food was restricted during fasting). All animal experiments were approved by The Roslin Institute’s Animal Users Committee and the animals were maintained in accordance with UK Home Office guidelines for the care and use of laboratory animals.

### Metabolic Studies

Male juvenile and adult (35 and 120 days old respectively) were weighed and fasted for 4 hours between 9am and 1pm. Prior to the start of the tests, a basal blood sample was collected by venesection into EDTA powder coated capillary tubes (Starstedt, Leicester, UK). Basal glucose levels were measured using a glucose monitoring system (Accu-Chek^®^ Aviva, Roche, Leicester, UK). 2 mg of D-glucose (Sigma, Poole, UK) per g of body weight was administered by gavage or 0.5mU of insulin (Actrapid, NovoNordisk, Bagsvaerd, Denmark) per g/body weight was administered intraperitoneally (i.p.). At precisely 15, 30, 60 and 120 minutes following administration, blood glucose was measured with an Accu-Chek^®^ Aviva glucose meter (Roche Diagnostics Ltd, Lewes, UK) and insulin was measured by ELISA (ChrystalChem, Chicago, IL, USA). Animals were allowed to recover for two weeks prior to euthanasia. Tissues were collected for protein, gene and histological analysis.

### Serum measurements

Serum samples were prepared from blood collected by heart puncture of CO_2_ culled mice in the fasted state. Total, carboxylated (GLA13-OCN), undercarboxylated (GLU13-OCN) and uncarboxylated (GLU-OCN) osteocalcin (69), adiponectin and leptin (CrystalChem) were quantified by ELISA.

### Primary osteoblast isolation and culture

Under sterile conditions calvaria were isolated from 2-4 day old new-born WT and *Phospho1^-/-^* mice as previously described (70). Osteoblasts were expanded in flasks in growth medium consisting of α-MEM supplemented with 10% FBS and 1% gentamicin in a humidified atmosphere of 95% air/5% CO_2_ and maintained at 37 °C. When the cells reached 80–90% confluency, they were seeded at a density of 2.5×10^4^/cm^2^ in multi-well plates. Conditioned medium was collected upon plate confluency, centrifuged to remove particulates and frozen at −80°C until required. For overexpression studies, primary osteoblasts were transfected with empty (EV) or overexpressing (OE) vectors as previously described (71).

### Mito Stress test

An XF24 Analyser (Agilent Technologies, Santa Clara, CA, USA) was used to measure the respiratory function of primary osteoblasts. Osteoblasts were plated at a density of 50,000 cells per well and transferred to a 37°C CO_2_ incubator until and calvarial osteoblasts were differentiated using osteogenic differentiation media containing 8 mM β-glycerophosphate and 50 μg/ml ascorbic acid for 3 days. On the day of the assay cells were washed in XF Assay Media supplemented with 25 mM glucose and 10 mM pyruvate and placed in a non-CO_2_ incubator at 37°C for 1 hour prior to start of assay. Reagents were prepared for the assay (injection volume of 75 μL for each reagent per well) from 2.5 mM Seahorse stock solutions, Oligomycin (1.2μM). Following equilibration the Seahorse plate was placed in the Seahorse XF24 Analyser for sample analysis. The raw data was normalized to protein concentration in each well at the end of the assay. The proton production rates and acidification correction were performed as described in (48).

### Gene expression analyses and immunoblotting

RNA extractions from tissues and cells was performed using the RNeasy Lipid Tissue Kit (Qiagen). The SuperScript First Strand Synthesis System (Invitrogen) was used for reverse transcription. Real-time PCR amplification with the 2 x precision master mix (Primer design, Southampton, UK) using the Stratagene Mx3000P real-time QPCR system (Agilent Technologies, Santa Clara, CA, USA). Each sample was tested in triplicate and compared to a housekeeping gene (Atp5B in osteoblasts and bone tissue, Lrp10 in adipose tissue and Gapdh in all other tissues) using MxPro software (Cheshire, UK) and the relative expression of the analysed genes was calculated using the ΔΔCT method (72). Primers sequences are available upon request. For protein extraction cells were scraped and tissues homogenised in an appropriate volume of radio-immunoprecipitation assay (RIPA) buffer containing 15% of complete mini protease inhibitor cocktail (Roche, Burgess Hill, West Sussex, UK). Protein concentration was determined by the Bio-Rad DC protein assay (Bio-Rad, Hertfordshire, UK). Immunoblotting was conducted with specific antibodies and protein bands were visualized using the enhanced chemiluminescence (ECL) Western Blotting Detection System (GE Healthcare, Chalfont St Giles, UK) or the Odyssey infrared detection system (LICOR).

### Tissue histology

Tissue was fixed in 4% PFA and embedded in paraffin wax. 5-μm sections were stained with hematoxylin and eosin (H&E) using the Leica Autostainer and mounted in DePeX (VWR, Lutterworth, UK). Adipocyte diameter and pancreatic β-cell islet number and size were quantified using ImageJ software as previously described (7, 73).

### Osmium staining

Mouse tibiae were fixed in 10% neutral-buffered formalin and decalcified in 14% EDTA, pH 7.4. Mouse bones were stained according to (74). Briefly, bones were stained with a 1% osmium tetroxide solution for 48 hours at room temperature. Bones were washed in Sorensen’s buffer and embedded in 1% agarose prior to μCT scanning (μCT100 Scanco Medical, Bassersdorf, Switzerland −12 μm, medium resolution, 70 kVp, 114 μA, 0.5 mm AL filter and integration time 500 ms. Analysis was performed using the manufacturer’s software.

### Micro-magnetic resonance, computed tomography and liver spectroscopy

CD and HFD mice were sacrificed immediately before imaging. For micro-magnetic resonance mice were imaged on a Varian 7 Tesla magnet using VnmrJ Pre-Clinical MRI Software. T2 weighted images were acquired both in the axial (1mm Thickness, 192×192 pixels, TR - 3000ms, TE 24ms, 1 average, FOV - 38.4 x38.4) and coronal planes (0.5mm Thickness, 512×256 pixels, TR - 3000ms, TE 24ms, 4 averages, FOV - 102.4×51.2). Cross-sectional computed tomography images were taken along the length of the body (3 mm apart, field of view 450 mm, approximately 70 images per mouse). Sheep Tomogram Analysis Routines (STAR) software (BioSS - V.4.8; STAR: Sheep Tomogram Analysis Routines, University of Edinburgh, http://www.ed.ac.uk) was used to calculate the total area and average densities of fat, muscle and bone in each carcass image without gutting (segmenting out guts and organs), based on density thresholds (low fat: −174 HU, high fat: −12 HU, low muscle:-10 HU, high muscle: 92 HU, bone: < 94 HU) (75).

Liver spectroscopy was conducted on user defined areas (TR - 1800ms, TE - 11.5 ms, 16 averages, Vauxhall 3×3×3). Lorentzian and Gaussian lineshape were used to fit peaks to MR data (jMRUI http://www.mrui.uab.es/mrui/mrui).

### Microarray and pathway analysis

Labelled cRNA was prepared from 500 ng of WT and *Phospho1^-/-^* primary calvarial osteoblast RNA using the Illumina^®^ RNA Amplification Kit from Ambion (Austin, TX, USA). The labeled cRNA (1500ng for mouse and 750ng for human) was hybridized overnight at 58°C to the SentrixMouseWG-6 Expression BeadChip or humanHT-12 Expression BeadChip (>46,000 gene transcripts; Illumina, San Diego, CA, USA) according to the manufacturer’s instructions. BeadChips were subsequently washed and developed with fluorolink streptavidin-Cy3 (GE Healthcare). BeadChips were scanned with an Illumina BeadArray Reader. Data was generated from Imagedata using Illumina software, GenomeStudio. Normalized data was generated using ^3^Cubic Spline^2^ Model in software. Pathway analysis was performed with Ingenuity Pathway Analysis (IPA, Ingenuity^®^ Systems, www.ingenuity.com) and GeneMANIA (http://www.genemania.org).

### Proteomic analysis

Proteins from serum was be extracted and prepared as previously described (76–78) and the extracted peptides was analysed using a RSLC 3000 nanoscale capillary LC followed by qTOF mass spectrometry (5600 Triple-TOF, Sciex). Sequential window acquisition of all theoretical spectra (SWATH) was used to profile all proteins in each sample using a data-independent acquisition method (79). ProteinPilot™ was used for protein identification and quantitation, as well as visualising peptide-protein associations and relationships.

### Choline Extraction

Serum samples were analysed using tandem mass spectrometry (LC-MS/MS) and a multiple-reaction monitoring (MRM) methodology. Five microliters of serum were extracted with 90 μL of an organic solution (10% methanol and 90% acetonitrile), containing the deuterium-labeled internal standard (IS, D9-Cho at 10 μg/mL). This resulted in precipitation of proteins, which were removed by filtration with a Millex 0.45 μm filter followed by centrifugation for 2 min at 6000 *g.* The mass transitions used to measure the analytes are: choline (mass transition *m/z* 104 → 60) and D9-choline (mass transition *m/z* 113 → 69). A QTRAP 5500 triple-quadrupole mass spectrometer (AB Sciex, Warrington, Cheshire, UK) with ESI ion source was used for data acquisition. Separation of analytes was performed in an Acquity UPLC-MS/MS (Waters, Hertfordshire, UK), with a binary pump system at a flow rate of 0.3 mL/min, connected to the mass spectrometer. The injection volume was 10 μL. Samples were separated using a Cogent 100mm×2.1 mm, 4 μm Diamond Hydride silica column (Microsolv Technologies, NJ, USA) and a linear gradient from 65% buffer B (0.1% formic acid in Acetronitrile) and 35% buffer A (0.1% formic acid in water) to 35% buffer B over 7 minutes. Analyst software (AB SCIEX) was used for HPLC system control, data acquisition, and data processing.

### Statistics

The data were analysed using various statistical models. All data were checked for normality and equal variance. Linear regression and correlation analysis based on Excel (Microsoft Office 10) built-in functions with interval of confidence and testing of the correlation coefficients were performed according to standard procedures. The SAS software was used to fit the generalized linear model (Microsoft Office 10). The Student’s *t*-test, ANOVA and a Two Way Repeated Measures ANOVA (Two Factor Repetition) for normally distributed data and non-parametric data was analysed using a Mann–Whitney Rank Sum test using Sigma Plot software (v 11.0) Systat Software Inc., London, UK) and Prism software (GraphPad, USA).Data are presented as means ± standard error (SEM) were appropriate. Regression and correlation coefficient’s are given with the intervals of confidence (p=0.05).

## Acknowledgments

We thank Elaine Seawright for technical assistance; Darren Smith for providing animal support; Dr Calum Gray, Prof. Maurits Jansen and Ross Lennen for MRI assistance (Edinburgh Imaging, University of Edinburgh); Prof. Roland. Stimson (Centre for Cardiovascular Science, University Edinburgh) for critical feedback on this manuscript. This project was funded by the Biotechnology and Biological Sciences Research Council (BBSRC) UK through a studentship award (KJS), and Institute Strategic Programme Grant Funding (BB/J004316/1 and BB/P013732/1) (CF, VEM), BBSRC Institute Career Path Fellowship funding from the BBSRC (VEM) and grant AR53102 from the National Institute of Arthritis and Musculoskeletal Diseases (NIAMS) from the National Institutes of Health (NIH), USA. Small project grants and lab exchange funds were provided by the Roslin Institute and The Society for Endocrinology.

