## Supplemental data 1 for "PHOSPHO1, a novel skeletal regulator of insulin resistance and obesity"

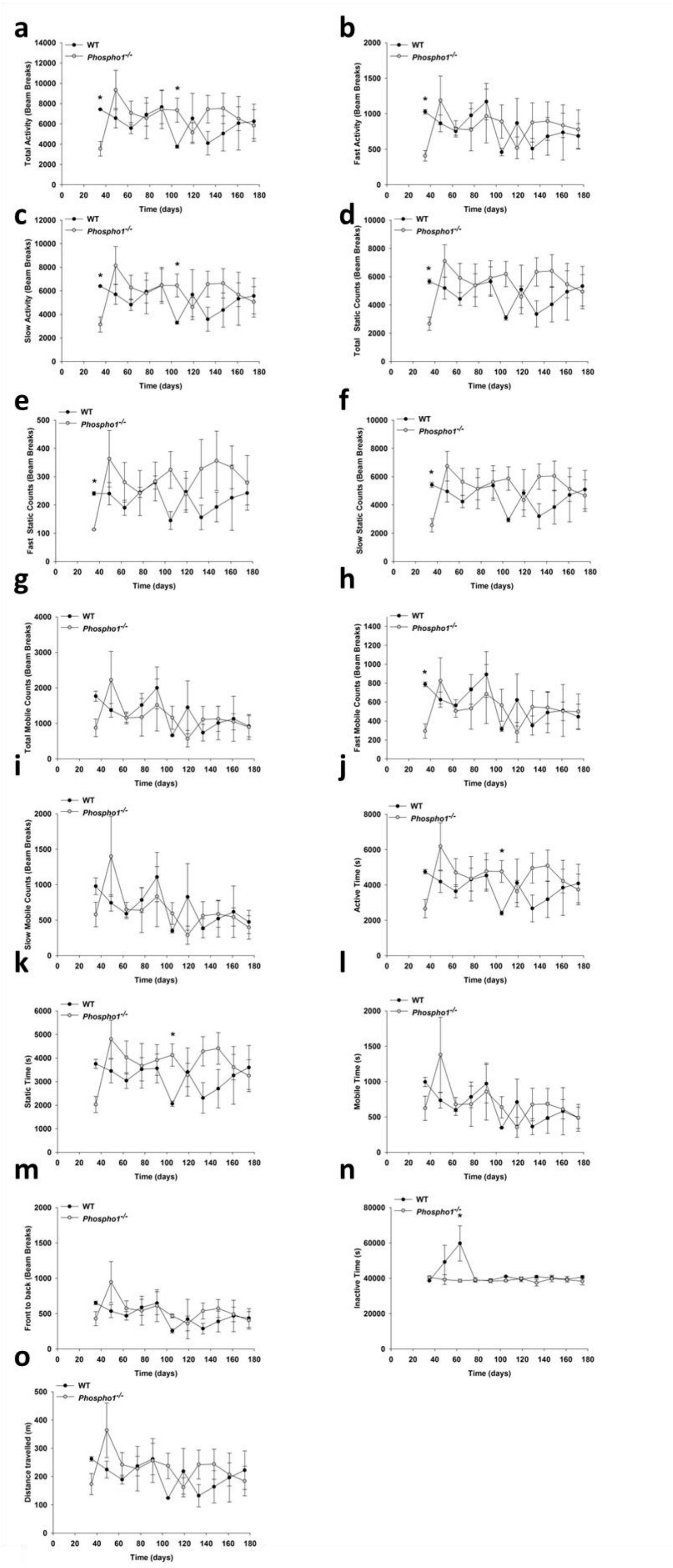


**Supplementary 1. Ambulatory activity of WT and *Phospho1*$^{-/-}$ mice**

**(a)** Total activity, **(b)** fast activity, **(c)** slow activity, **(d)** total static counts, **(e)** fast static counts, **(f)** slow static counts, **(g)** total mobile counts, **(h)** fast mobile counts, **(i)** slow mobile counts, **(j)** active time, **(k)** static time **(l)** mobile time, **(m)** front to back, **(n)** inactive time, **(o)** distance travelled. Data are represented as mean ±S.E.M (n=6 replicates). *$p<0.05$.
