## Supplemental data 2 for "PHOSPHO1, a novel skeletal regulator of insulin resistance and obesity"

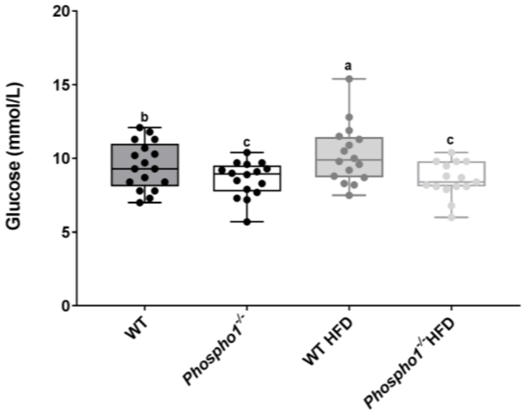


**Supplementary 2.** Fasted glucose levels of 120 day old WT and *Phospho1*$^{-/-}$ mice on the control and HFD. Different letters above the error bar for each gene show significant difference at $p<0.05$.
