## Supplemental data 3 for "PHOSPHO1, a novel skeletal regulator of insulin resistance and obesity"

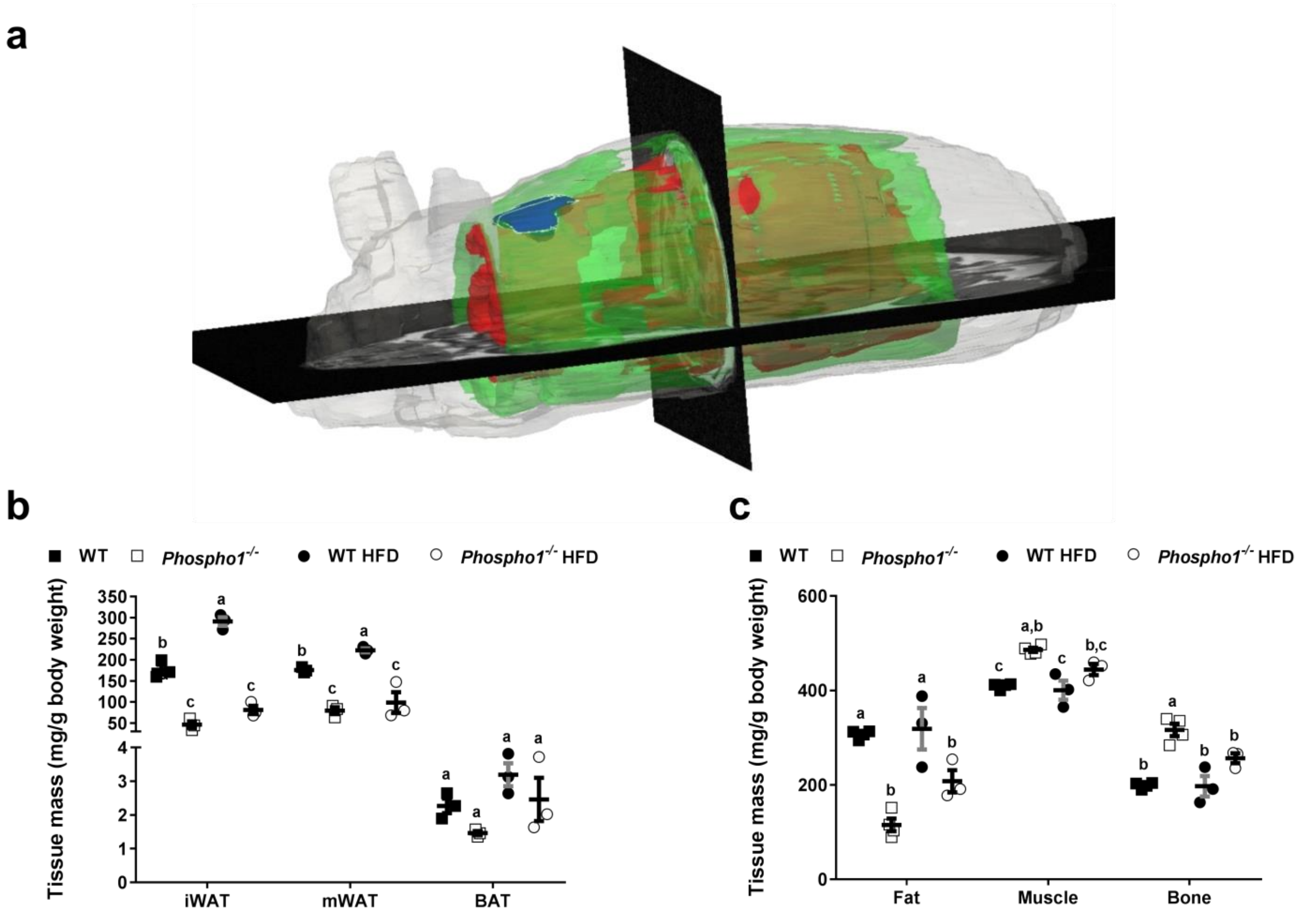


**Supplementary 3. µMRI and CT fat quantification from WT and *Phospho1*$^{-/-}$ mice on both a control and HFD. (a)** Representative reconstructed µMRI scan. Green = subcutaneous adipose tissue, Red = mesenteric adipose tissue, Blue = brown adipose tissue. (**b)** Inguinal WAT (iWAT), mesenteric WAT (mWAT) and brown adipose tissue (BAT) mass determined by µMRI. **(c)** Fat, muscle and bone mass determined by non-segmented multi-object CT scanning. Results were normalised to body weight (mg/g). Data are represented as mean ±S.E.M (n=6 replicates). Different letters above the error bar for each gene show significant difference at $p<0.05$.
