## Supplemental data 4 for "PHOSPHO1, a novel skeletal regulator of insulin resistance and obesity"

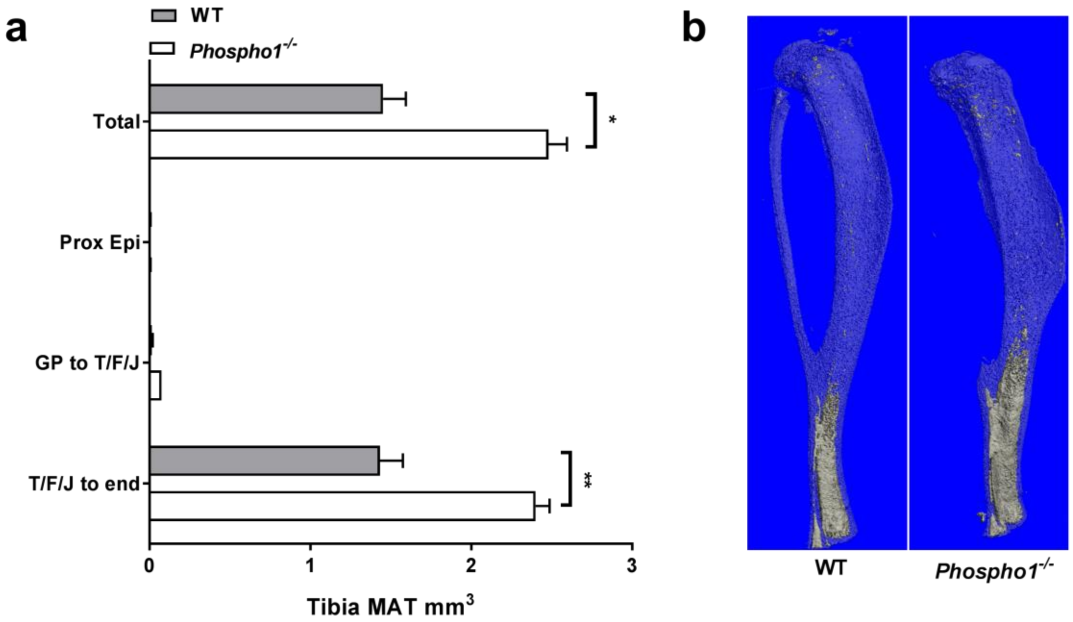


**Supplementary 4. Marrow adipose tissue µCT osmium quantification. (a)** Region-specific quantification of tibial marrow adipose tissue (MAT) volume. Regions include the proximal epiphysis (Prox Epi), the growth plate to the tibia/fibula (Tib/Fib) junction (GP to T/F J) and the tibia/fibula junction to the end of the bone (T/F J to end). **(b)** Representative images of osmium-stained tibiae scanned by µCT. Marrow fat is dark grey and bone is light grey Data are represented as mean ±S.E.M (n=3 replicates). * $p<0.05$, ** $p<0.01$.
