## Supplemental data 5 for "PHOSPHO1, a novel skeletal regulator of insulin resistance and obesity"

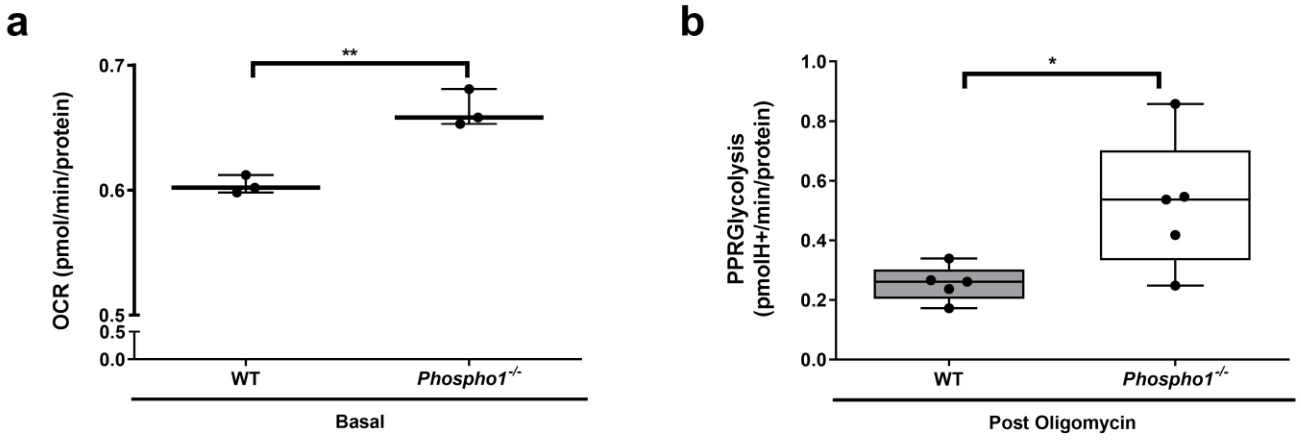


**Supplementary 5. Seahorse analysis of WT and *Phospho1*$^{-/-}$ osteoblasts. (a - b)** Oxygen consumption rates and proton production rates (calculated from ECAR) using Seahorse X-24 analyzer in WT and *Phospho1*$^{-/-}$ primary calvarial osteoblasts under basal conditions (OCR) or following the addition of oligomycin (PPRGlycolysis). Data are represented as means ±S.E.M from five individual wells for each group and is representative of two independent seahorse runs. * $p<0.05$, ** $p<0.01$.
