## Supplemental data 6 for "PHOSPHO1, a novel skeletal regulator of insulin resistance and obesity"

| **Gene in Dataset** | **Fold From WT** |
|---|---|
| ABHD16A | -14.224 |
| ALOX5AP | -3.427 |
| Ccl9 | -3.863 |
| CLEC6A | -3.787 |
| CTSS | -2.372 |
| FCER1G | -15.853 |
| FCGR2A | -3.299 |
| FCGR2B | -2.432 |
| Gp49a/Lilrb4 | -3.138 |
| Gp49a/Lilrb4 | -3.138 |
| GPNMB | -13.704 |
| HLA-B | -18.680 |
| ICAM1 | 2.527 |
| Ifi202b | 11.328 |
| Lyz1/Lyz2 | -11.330 |
| MPEG1 | -7.108 |
| MRC1 | -2.510 |
| MS4A6A | -4.583 |
| SERPING1 | 1.993 |
| TYROBP | -2.971 |
| VDR | 2.297 |
| VIP | 2.093 |

**Supplementary 6. Osteoblast microarray candidates involved associated with glucose homeostasis.**
21 genes from the WT and *Phospho1*$^{-/-}$ osteoblast microarray were identified by Ingenuity Pathway Analysis to be associated with glucose homeostasis p=1.04x10$^{-6}$.
