## Supplemental data 7 for "PHOSPHO1, a novel skeletal regulator of insulin resistance and obesity"

| Gene | Function in Diabetes | Function in Bone | Ingenuity Prediction | Fold From WT | Fold | p value | * rating |
|---|---|---|---|---|---|---|---|
| ***Vdr*** | Deficiency associated with T1DM. | The active form, 1alpha,25-(OH)2D, binds to the vitamin D receptor (VDR) to modulate gene transcription and regulate mineral ion homeostasis. | ↑ | ↑ | 2.21 | 0.00017 | *** |
| ***Mpeg1*** | WAT-associated inflammation in (HFD) is accompanied by reduced expression of immune and inflammatory response genes including MPEG1 | Correlated with correlation femur ultimate force. | ↓ | ↑ | 3.93 | 0.00011 | *** |
| ***Slc1a3 (glutamate transporter)*** | Alterations in glutamate transport during diabetes (retina) | | ↑ | ↑ | 4.16 | 0.00003 | *** |
| ***Adamts4*** | ADAMTS families are regulated by PPARγ, which increases insulin sensitivity. | Key enzymes involved in the cleavage of aggrecan and the degradation of cartilage | ↑ | ↑ | 1.67 | 0.00164 | ** |
| ***Bmp4*** | BMP4 has been suggested to play an important role in adipogenesis, especially the white adipocyte differentiation through interaction with BMP receptor (BMPR) and subsequently activating the Smad signaling pathways | BMP-4 stimulates osteocalcin synthesis in osteoblast-like MC3T3-E1 cells. | ↓ | ↓ | 0.12 | 0.00004 | *** |
| ***Cd68 (macrophage transmembrane protein)*** | Oobesity is associated with significant infiltration of adipose tissue by macrophages.<br><br>Treatment with pioglitazone reduces expression of CD68 and MCP-1 in adipose tissue, apparently by reducing macrophage numbers, resulting in reduced inflammatory cytokine production and improvement in insulin sensitivity. Higher CD68 mRNA levels with obesity and insulin resistance. | Genetic ablation of CD68 results in mice with increased bone and dysfunctional osteoclasts. | ↓ | ↓ | 0.56 | 0.01518 | * |
| ***Cfp (complement factor properdin)*** | Elevated Properdin and Enhanced Complement Activation in First-Degree Relatives of South Asian Subjects With Type 2 Diabetes | Complement proteins are present in developing endochondral bone and may mediate cartilage cell death and vascularization - is localized in the resting and hypertrophic zone but not in the proliferating zone is localized in the resting and hypertrophic zone but not in the proliferating zone. | ↓ | ↓ | 0.24 | 0.00002 | *** |
| ***Cxcl4*** | Incubation of 3T3-L1 adipocytes with CXCL14 stimulated insulin-dependent glucose uptake CXCL14 plays a causal role in high-fat diet-induced obesity | ex vivo expansion of HSCs may be highly effective through osteoblast-differentiated MSCs acting as a feeder layer, and likely operates through the CXCL12 chemokines signalling pathway. | ↑ | ↑ | 3.51 | 0.00002 | *** |
| ***Fmod*** | Fibromodulin were found to be downregulated after 40 weeks of diabetes. | Important in maintaining periodontal homeostasis through regulation of TGFβ/BMP signalling, matrix turnover, and collagen organization. | ↑ | ↑ | 1.34 | 0.00031 | *** |
| ***Lum (lumican)*** | Collagen-associated proteoglycan. In the present study, increased deposition of collagen types I and V and decreased deposition of collagen type III, biglycan and lumican was observed in the decidua of the diabetic group (uterine lining (endometrium) | lumican is a significant proteoglycan component of bone matrix, which is secreted by differentiating and mature osteoblasts only and therefore it can be used as a marker to distinguish proliferating pre-osteoblasts from the differentiating osteoblasts. | ↑ | ↑ | 6.80 | 0.00000 | *** |

**Supplementary 7. *In Silico* analysis of Ingenuity pathways predictions** *In Silico* analysis of genes predicated to be associated with bone and diabetes mellitus.
