## Supplemental data 8 for "PHOSPHO1, a novel skeletal regulator of insulin resistance and obesity"

| Unique serum proteins | Molecules in Network | Score | Focus Molecules | Top Diseases and Functions |
|---|---|---|---|---|
| WT HFD | ACE,Akt,Alpha catenin,APOM,CD3,COL1A1,Collagen type I,Collagen type IV,Collagen(s),CTSB,ERK,ERK1/2,F7,F9,HNF1A,IHH,IL1,Insulin,LDL,LYZ,MST1,P38 MAPK,Ph,PI3K (complex),PLA2G7,POSTN,PROS1,PSAP,PSMA5,QPCT,TCF,TFRC,Tgf beta,TGFBI,Vegf | 47 | 18 | Hematological System Development and Function, Organismal Functions, Connective Tissue Disorders |
| WT HFD | CSNK1G3,EFEMP1,FIG4,KRT75,MTMR4,MTMR6,MTMR8,MTMR9,NT5C,NUDT11,PDXP,PGAM4,PPAPDC2,PPIP5K1,PPP1R14A,PPP2R3B,PPTC7,PTPLB,RNF166,SACM1L,SSH3,THTPA,TPTE,UBC,UBLCP1,UFL1,USP30,USP34,USP35,USP38,USP40,USP44,USP48,USP27X,USP9Y | 12 | 6 | Post-Translational Modification, Carbohydrate Metabolism, Lipid Metabolism |
| WT HFD | LHX1,SPP2 | 3 | 1 | Developmental Disorder, Embryonic Development, Organ Development |
| WT HFD | HNRNPA2B1,MAST4,SMAD1 | 2 | 1 | Tissue Development, Connective Tissue Disorders, Developmental Disorder |
| WT HFD | ARHGAP20,DMD,ERG,FIGLA,Rap1,SRF,TERF2 | 2 | 1 | Cellular Function and Maintenance, Cell Cycle, Reproductive System Development and Function |
| WT HFD | ARHGAP18,BMP6,CDKN2A,MPHOSPH6,MPP6,RB1,RHOA,WNT3A | 2 | 1 | Cancer, Tumor Morphology, Cellular Growth and Proliferation |
| WT HFD | C1orf94,DMRTB1,EWSR1,EYA2,IL16,LENG8,NCKIPSD,NEDD9,NIF3L1,PEF1,PRMT2,RBFOX1,RBFOX2,SORBS3,VASP | 2 | 1 | Cell-To-Cell Signalling and Interaction, Nervous System Development and Function, Cellular Assembly and Organization |
| *Phospho1*$^{-/-}$ HFD | ACO1,aldo,ALDOB,ASL,Beta Tubulin,BHMT,CAT,DBI,ELF1,ENO1,ERK1/2,FABP1,FBP1,GAPDH,glutathione peroxidase,glutathione transferase,GPX1,GSTA3,GSTM5,GSTZ1,HNF4α dimer,Kap,Ldh (complex),MAT1A,MDH1,PARK7,PF4,PGAM1,PGK1,PRDX6,PROC,SELENBP1,SLC4A1,Sod,SOD1 | 60 | 27 | Carbohydrate Metabolism, Free Radical Scavenging, Small Molecule Biochemistry |
| *Phospho1*$^{-/-}$ HFD | 14-3-3,ADH4,ADH5,ADH1C,ADK,AHCY,Akt,alcohol dehydrogenase,Alpha tubulin,APCS,ASS1,caspase,CCT5,cytochrome C,EEF2,EEF1A1,FAM188B,GLO1,GLUL,HSP,HSP90AB1,HSPA8,LDHA,MTHFD1,MYLK2,PRDX1,PRKDC,PSMA1,SEC14L2,SPTA1,TCR,TUBB4B,VCAM1,YWHAE,YWHAZ | 60 | 27 | Energy Production, Small Molecule Biochemistry, Drug Metabolism |
| *Phospho1*$^{-/-}$ HFD | CHKB,CNDP2,DAK,DCAKD,DCXR,ECT2,EED,EPHX2,ESD,ETNK1,KRT5,LAP3,MB21D2,MFSD10,MTMR8,NAA40,NAV3,NRDE2,NRM,NSUN5,OBSCN,OSTC,PGAM4,PGM1,PLEKHG3,PPWD1,RBM12B,SORD,SZT2,TBC1D31,TEKT2,THTPA,TSG101,UBC,VPS37D | 28 | 15 | Carbohydrate Metabolism, Lipid Metabolism, Small Molecule Biochemistry |
| *Phospho1*$^{-/-}$ HFD | AKR1C4,ALDH,ALDH1A1,ALDH1L1,ALDH8A1,AMPK,Ap1,ATP9A,CD3,CD5L,Creb,FSH,Histone h4,HPD,HRSP12,IgG,Iglv1,IL12 (complex),Immunoglobulin,LDL,Mapk,MIB2,MSRA,NADPH oxidase,NFkB (complex),NME1,P38 MAPK,PI3K (complex),PPIA,Pro-inflammatory Cytokine,PYGL,retinal dehydrogenase,SRC (family),Tgf beta,Vegf | 25 | 14 | Drug Metabolism, Lipid Metabolism, Small Molecule Biochemistry |
| *Phospho1*$^{-/-}$ HFD | Ahsp,ANKS1A,APOL1,AQP11,ARL6,C10orf54,CA1,CD44,CELA1,CLEC5A,Cma2/Mcpt9,Cyb5r3,DOK5,EGFR,EPO,F2,F3-F7,FAH,FGFRL1,FTCD,HAAO,HOOK2,HYAL2,IFNG,NCF1C,NLRP10,P4HTM,PAH,PBLD,PCTP,Podxl,PPARA,SERPINB10,SLC46A1 | 12 | 10 | Cell Signalling, Small Molecule Biochemistry, Vitamin and Mineral Metabolism |
| *Phospho1*$^{-/-}$ HFD | 26s Proteasome,Actin,Akt-Calmodulin-Hsp90-Nos3,ARG1,CA3,Calmodulin,Calmodulin-Hsp90-Nos3,CAP2,Cg,Ck2,Collagen type IV,CORO6,ERK,ERMAP,FAM3D,FKBP51-TEBP-GR-HSP90-HSP70,Focal adhesion kinase,GNMT,Histone h3,Hsp70,Hsp90,Insulin,Integrin,Jnk,KIR2DS4 (includes others),Lh,MUC8,p85 (pik3r),PIH1D3,Pkc(s),RGN,RNA polymerase II,THBS4,TINAG,TROPONIN | 7 | 5 | Amino Acid Metabolism, Embryonic Development, Organismal Development |

**Supplementary 8. *In Silico* analysis top diseases associated with unique WT HFD and *Phospho1*$^{-/-}$ HFD proteins.**
