## Supplemental data 9 for "PHOSPHO1, a novel skeletal regulator of insulin resistance and obesity"

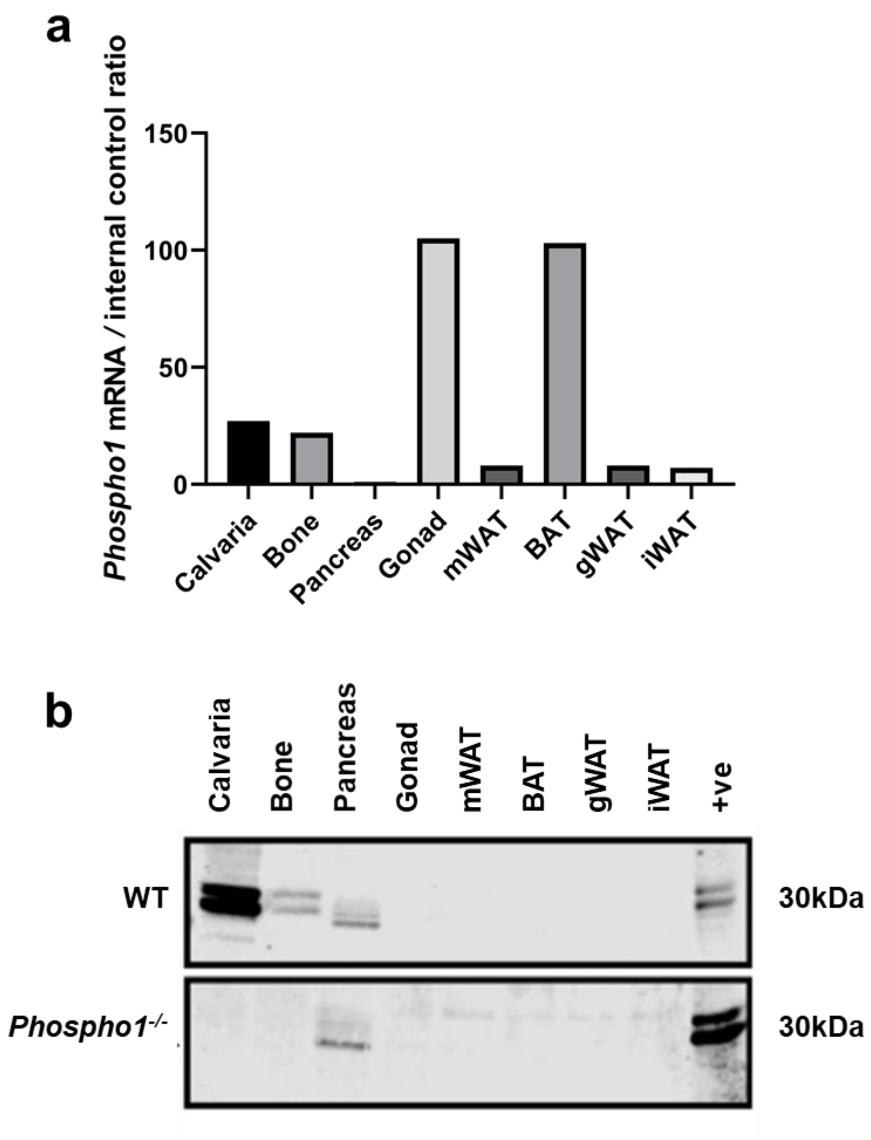


**Supplementary 9. Gene and protein expression of *Phospho1* mRNA and PHOSPHO1 protein in murine tissue**
**(a)** RT-qPCR of *Phospho1* in murine tissues, high expression was seen in the gonad and brown adipose tissue (BAT) (**b**) Protein expression of PHOSPHO1 was detectable by western blot in the calvaria and bone. Non-specific binding of the PHOSPHO1 antibody was observed in the pancreas, seen in both WT and *Phospho1*$^{-/-}$ pancreatic tissue.
